# LightSuite: scalable atlas registration and cell detection across the central nervous system

**DOI:** 10.64898/2026.10.13.757638

**Authors:** Dimokratis Karamanlis, Yuqiao Xie, Charlie G. Foucher, Melvin Alappat, Anas Masood, Ioanna Dimitropoulou, Louis Ange Félix Pivron, José María Martínez de Paz, Emilie Macé, Sevasti Gaspari, Sami El-Boustani

## Abstract

Deciphering brain-wide circuits requires large animal cohorts, yet computational bottlenecks in atlas registration and whole-volume cell detection reduce precision, reproducibility, and sample yield. We present LightSuite, an open-source framework enabling scalable, multimodal registration and cellular mapping across the central nervous system (CNS). LightSuite combines damage-resilient 3D point-cloud initialization with user-steerable non-rigid registration, and GPU-accelerated local background normalization for rapid, illumination-invariant cell detection. Applying LightSuite to Fos-dependent activity labeling in mice, we extracted reproducible, layer-resolved profiles of active cell density, soma-size distributions and per-animal labeling-efficiency factors. LightSuite computationally untwists and registers cleared spinal cords to map myeloarchitecture, reconstructs serial wide-field histology to map surface GluA1 distributions, resolves optic fiber tracks, and aligns *in vivo* functional ultrasound hemodynamics into common atlas coordinates. LightSuite establishes an integrated, accessible platform for cohort-level structural, molecular, and functional CNS mapping.

## MAIN

Tissue clearing^1–4^ and light-sheet microscopy^5–8^, combined with activity-dependent genetic tagging^9,10^, enable brain-wide cellular mapping of behaviorally and pharmacologically relevant circuits^11–15^. At cellular to subcellular resolution, a single cleared brain can readily reach the terabyte scale, capturing the three-dimensional position and morphology of every labeled cell. Yet because each animal provides only a single activity snapshot, reproducible inference with respect to the ongoing behavioral or pharmacological state demands large animal cohorts. This requirement thus gives rise to a computational bottleneck: precisely registering each sample to a reference coordinate atlas^16^ and segmenting individual cells at whole-volume scale on standard laboratory hardware.

Existing pipelines fail to combine damage-resilient registration with scalable cell detection across modalities. Automated registration tools relying solely on global volume similarity^17–19^ frequently fail on distorted or incomplete tissues, leading to a substantial loss of usable samples and compromised experimental yield. Furthermore, standard frameworks lack intuitive interfaces to systematically incorporate user-guided anatomical priors^20^. Registering *in vivo* functional imaging modalities is harder still. In functional ultrasound^21,22^ (fUS), vascular contrast deviates markedly from cytoarchitectural reference atlases, complicating comparisons of functional responses across brains. On the cellular scale, deep-learning segmentation networks^23–25^ achieve high accuracy within small volumes, but whole-brain deployment requires high-memory computing clusters^11,26^ and remains slow. Finally, current workflows do not generalize beyond the intact brain to the spinal cord or other organs, nor do they recover the three-dimensional trajectories of implanted optical fibers to evaluate the targeting precision of deep brain imaging^27^ or optogenetic manipulations^28^.

To address these challenges, we developed LightSuite, an open-source framework for scalable atlas registration and cell detection across the central nervous system. LightSuite combines damage-tolerant 3D point-cloud initialization with an active-learning interface that directly embeds user-defined anatomical landmarks into a multi-metric non-rigid registration objective. For cellular quantification, it uses GPU-accelerated local signal-to-background normalization coupled with a lightweight convolutional artifact classifier, enabling fast, illumination-invariant soma detection on standard workstations. Each module runs independently or as an end-to-end workflow. We benchmarked LightSuite by extracting reproducible, layer-resolved profiles of Fos-dependent tdTomato-expressing cell density and soma size in TRAP2×Ai14 mice^9^. Across other CNS preparations and imaging modalities, LightSuite computationally straightens and registers cleared spinal cords, reconstructs serial wide-field histological sections, resolves 3D fiber tracks, and aligns *in vivo* power-Doppler fUS volumes into the Allen Mouse Brain Common Coordinate Framework^16^ (CCFv3).

## RESULTS

### A pipeline for whole-brain analysis

Mapping neural networks that relate to a specific behavior demands increasingly larger cohorts of animals, much as reproducible inference in human functional magnetic resonance imaging requires large sample sizes^29^. Rodent cohorts are costly: behavioral paradigms and clearing protocols are time-consuming and resource-intensive, so discarding a sample is expensive. Yet samples that show deformation, variable damages or missing structures^30^ often defeat existing analysis pipelines^19,31,32^. LightSuite standardizes registration and cell detection across heterogeneous whole-brain volumes end to end (Fig. 1), so that imperfect samples can be retained rather than discarded.

**Fig. 1.**
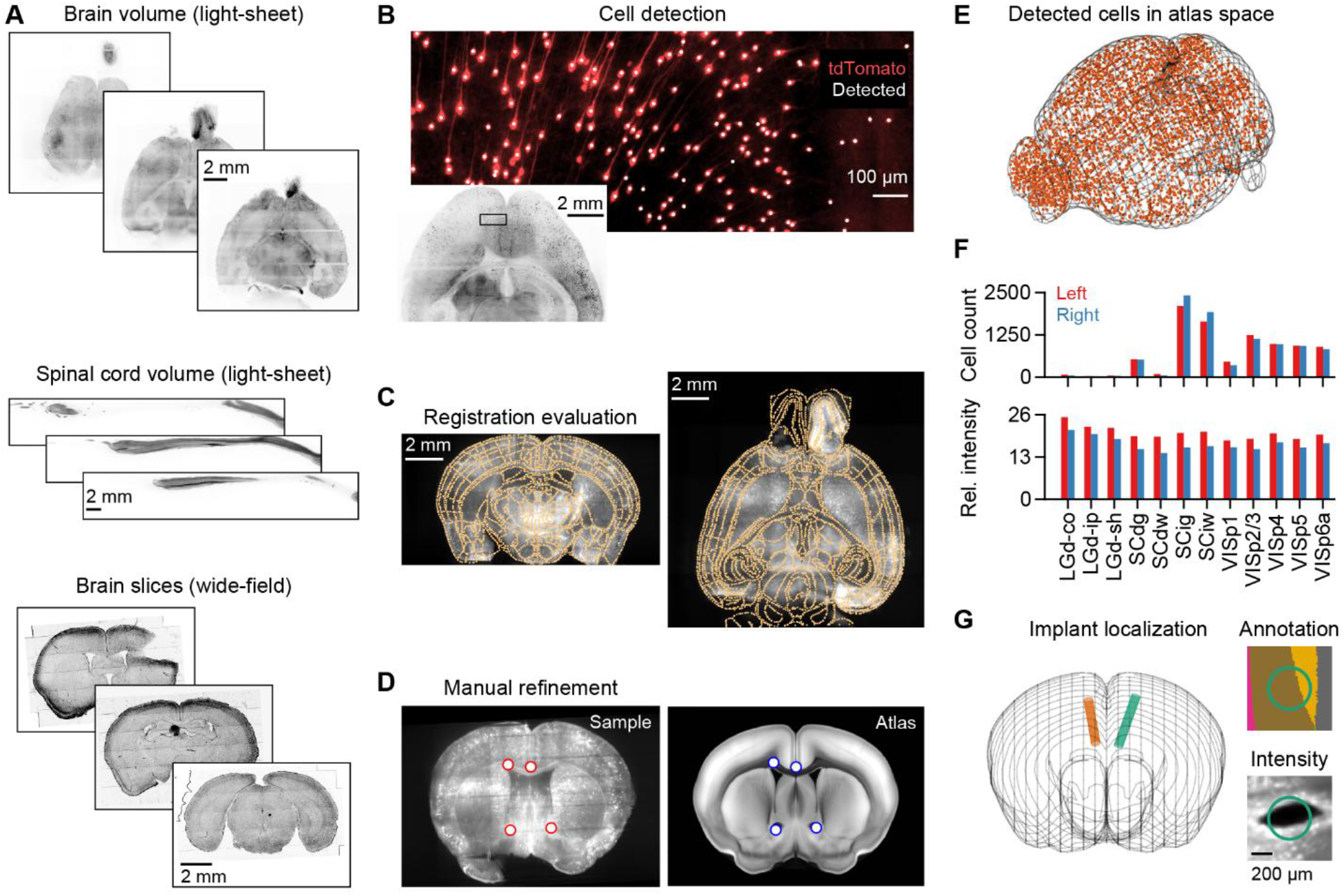
Overview of LightSuite. **A**) Three example slices for each supported sample type; from top to bottom: brain volume (light-sheet), spinal cord (light-sheet), and serially sectioned brain slices (wide-field images of coronal slices stained with DAPI). **B**) Example cell detection results in a small brain patch; detected cells (white dots) are overlaid on a subvolume with tdTomato-labeled neurons. Inset shows the location of the patch in the brain. **C**) Registration to the Allen Mouse Brain Atlas can be evaluated visually using atlas boundaries (gold) overlaid on two slices of an aligned sample (left: coronal; right: axial). **D**) Registration refinement via user-defined corresponding points mapped between approximately aligned 2D slices (left: sample; right: atlas). **E**) Detected cells in atlas space. **F**) Cell counts and relative fluorescence intensity for the left and right hemispheres across a subset of atlas substructures in the mouse visual pathway. Note the hemispheric asymmetry in intensity caused by unilateral light-sheet illumination. **G**) Localization of cylindrical implants (e.g., optic fibers) in atlas space (left), together with the estimated field of view at the fiber tip, both in atlas annotation space (right-top) and sample intensity space (right-bottom).

LightSuite can handle several sample types for which reference atlases exist (Fig. 1A). Cell detection runs at native resolution by extracting 3D maxima (Fig. 1B). For efficient registration, samples are then downsampled to 20-μm isotropic resolution, much finer than the non-rigid transform scale (see Methods), to be aligned to the atlas^16^ through a sequence of similarity, affine, and non-rigid transformations. Users can refine the latter two stages through a dedicated interface that introduces landmark points on slices already approximately aligned to the atlas (Fig. 1C-D). Outputs can include cell locations in atlas space (Fig. 1E), per-region cell counts and intensities for each hemisphere (Fig. 1F), as well as time-resolved regional signals for fUS.

LightSuite also reconstructs cylindrical fiber trajectories to evaluate targeting precision in optogenetic and imaging experiments. Users annotate the damage left by the implant track across serial slices (Ext. Data Fig. 1A), and the framework fits a 3D cylinder constrained by the local brain surface normal vector to determine the precise implant axis and terminal fiber face (Fig. 1G, Ext. Data Fig. 1B,C). This spatial readout directly resolves whether opsin or indicator expression falls within the effective photon cone, and can be used to assign cells within a field of view to different atlas regions.

To demonstrate LightSuite at cohort scale, we analyzed brain-wide fluorescent protein expression related to an immediate-early gene in TRAP2×Ai14 mice (Fos2A-iCreER×Ai14). In this model^9^, 4-hydroxytamoxifen (4-OHT) injection after a behavioral experience drives permanent, Cre-dependent tdTomato expression in active neurons, converting transient Fos expression into stable fluorescence. Throughout, we refer to the measured signal as tdTomato fluorescence and to the detected somata as tdTomato+ cells; both report Fos expression during the labeling window rather than the c-Fos protein itself.

### Point-cloud-assisted registration

Non-rigid registration of soft tissue is nonconvex and sensitive to initialization and degrees of freedom. LightSuite parameterizes the non-rigid transform with a B-spline grid spaced at several hundred micrometers (640 μm for mouse brains). This coarse grid spacing limits free parameters and prevents overfitting without a displacement-penalty regularizer, which adds computational cost in other pipelines^11,17,26^. This spatial scale reflects the physical mechanics of tissue preparation: fixation and clearing deform tissue smoothly^2,33,34^, producing displacements spanning several hundred micrometers to ∼1 mm. It also matches the natural delineation of the mouse brain, whose substructures and structures span several hundred micrometers (Ext. Data Fig. 2).

To initialize registration robustly, we first extracted points of high saliency at the spatial scale of brain structures, computed locally to control for inhomogeneous illumination (Fig. 2A,B). We assembled these points into a 3D structural point cloud in sample space and repeated the procedure for the atlas template volume. To align the two clouds, we fitted a similarity transform (rotation, translation, uniform scaling) to the atlas by Bayesian coherent point drift^35^, a point-cloud registration algorithm (Fig. 2C). A similarity rather than a rigid transform accounts for isotropic volumetric shrinkage or swelling during clearing. Unlike voxel-intensity initialization, point-cloud alignment^36^ inherently tolerates structural imperfections: localized tears, compression, and bubble voids are treated as statistical outliers, while missing regions (such as detached olfactory bulbs or cerebellum) remove points without displacing remaining anatomy. This robust initialization helps place the sample within the capture range of the true anatomy, mitigating the risk of downstream non-rigid optimization becoming trapped in unphysical local minima.

**Fig. 2.**
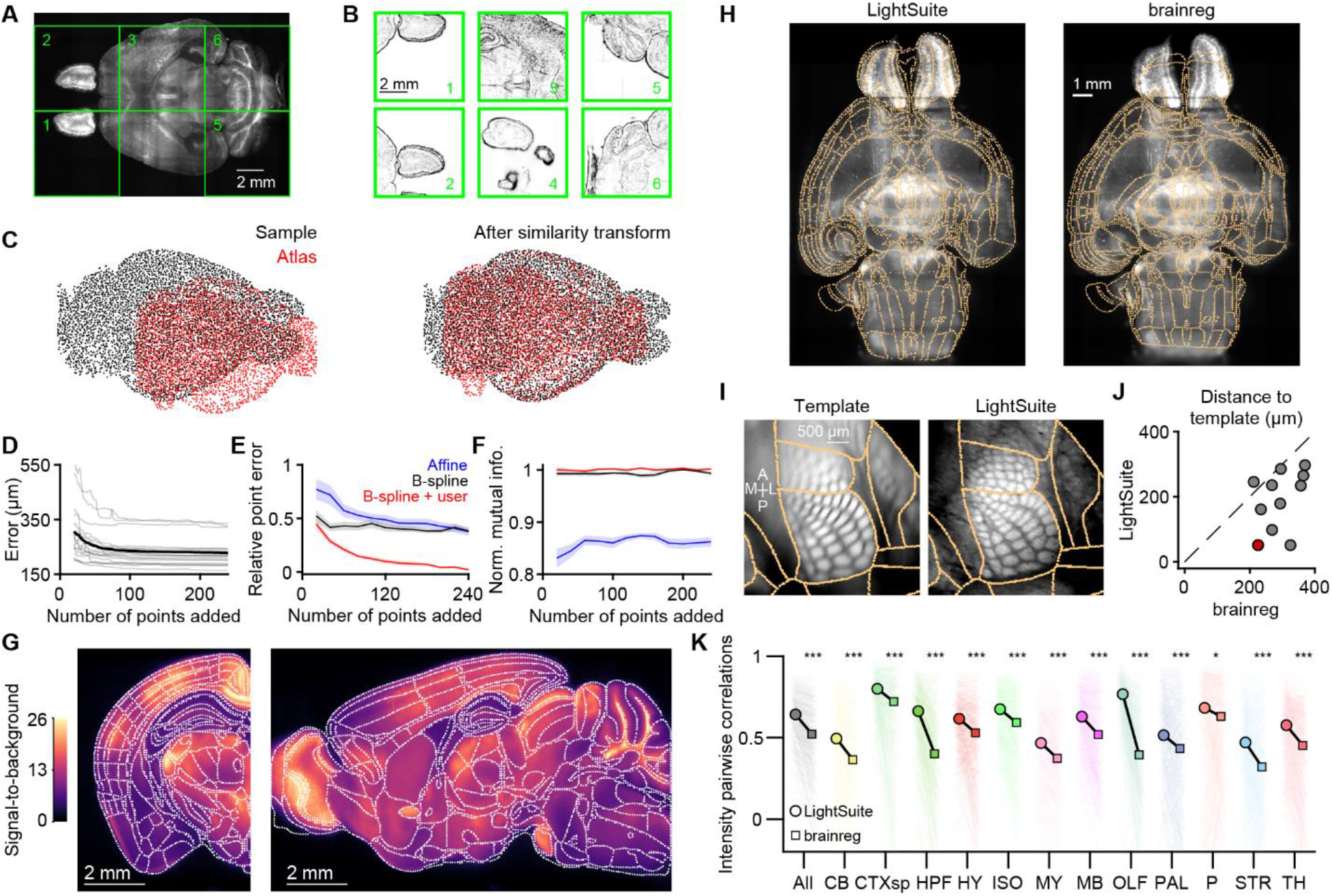
Consistent registration of cleared mouse brains with a mixed objective. **A)** Example axial slice of a light-sheet-acquired tdTomato-expressing volume. **B**) Six example local slice cutouts from the sample in panel A (green squares), used for automatically deriving point clouds after a gradient operation (see Methods). **C**) A coherent point drift algorithm fits a similarity transform to align sample (black, n=1.45·10^6^ points) and atlas (red, n=1.06·10^7^ points) point clouds (left: before; right: after transform). **D**) Root-mean-square error across manually placed user landmarks (excluding the initial automated point clouds) as a function of the number of landmarks added in chronological order. Individual samples (n=17) with available annotation order are shown in gray; mean is shown in black. **E**) Mean squared error relative to its value at the maximum number of landmarks for a B-spline transform combined with user landmarks (red). The same error metric is shown for landmark-free B-spline optimization (black; dependent on landmark count because it is initialized by the landmark-derived affine transform) and for the affine transform driven by landmarks alone (blue). Solid lines represent medians across samples (n=17); shaded regions indicate robust standard errors. **F**) Mutual information between sample and atlas volumes, normalized to the value attained by the B-spline fit with user landmarks at the maximum number of points added. Lines and shading as in (E). **G**) Average tdTomato fluorescence intensity (n=33 brains) shown in representative hemisected coronal (left) and sagittal (right) slices. Colormap indicates intensity relative to background outside the brain. White lines indicate Allen Mouse Brain Atlas structural boundaries. **H**) Example axial slice of a sample with overlaid area boundaries (yellow) registered with LightSuite (left) or brainreg (right). **I**) Left: Average flatmap projection of layer 4 of the Allen Mouse Brain Atlas reference template centered on the barrel cortex. Right: Corresponding projection of an autofluorescence sample (488 nm) registered with LightSuite. **J**) Comparison of barrel cortex displacement from the atlas template in flattened space between LightSuite and brainreg, with the sample from (I) highlighted in red. Values represent the root-mean-square displacement across the barrel field after 2D rigid alignment (see Methods). Displacement is lower for LightSuite than for brainreg (Wilcoxon signed-rank test, p=0.005, n=11 barrel fields from 9 brains; seven barrel fields could not be resolved by brainreg and were excluded). Flatmap distance provides a conservative estimate of true displacement, as brainreg-registered barrels frequently fell outside layer 4. **K**) Pairwise Spearman correlations of regional fluorescence intensity profiles (n=30 brains) analyzed with LightSuite (circles) and brainreg (squares) across Allen Mouse Brain Atlas substructures. LightSuite-registered samples exhibit higher pairwise correlations than brainreg-registered ones (n=435 pairs, Wilcoxon signed-rank test, p<10^-16^); this advantage holds for substructure intensities within each major brain division (*p<0.05, **p<0.01 and ***p<0.001).

Users steer the registration by adding corresponding anatomical landmarks between the sample and atlas template volumes: this sparse user-defined point cloud replaces the dense saliency clouds used for initialization. Because LightSuite inverts the initial similarity transform to bring the sample into atlas space, users place landmarks on orthogonally resliced coronal, sagittal and axial slices rather than searching for correspondences directly in 3D. An active-learning interface accelerates this annotation by fitting a running 3D affine transform from accumulated landmark pairs in real time. When a user marks an anatomical feature on a sample slice, the interface immediately suggests the matching 2D correspondence in the atlas view. The affine registration error, the root-mean-square distance across all landmarks, falls monotonically with the number of user points and saturates after roughly 100 landmarks (Fig. 2D). These landmarks carry local anatomical information absent in the global transform, and they enter the non-rigid fit directly: the B-spline transform optimizes a weighted combination of the mutual information between the sample and atlas volumes^34,37^ and the Euclidean distance between the paired points (see Methods). Adding the points sharpens the fit (Fig. 2E, Ext. Data Fig. 3). Mutual information exceeds the affine-only value but does not change with the number of landmarks, although it increases slightly when the landmarks enter the objective (Fig. 2F). These measurements expose a limitation of image-only registration: the landmark error keeps falling as anatomically informative points are added, while mutual information stays flat and so cannot separate an anatomically faithful alignment from an erroneous configuration that yields an equally high similarity score. The user landmarks supply the signal that mutual information misses, and their residual error is a more sensitive quality measure. Most landmarks we placed fell along ventricles and white-matter tracts (Ext. Data Fig. 4A,C), and annotation time scaled linearly with their number (Ext. Data Fig. 4B), giving precise registration within 10–20 minutes for each sample.

Standard light-sheet workflows acquire separate autofluorescence and signal volumes, doubling data acquisition time and storage. In contrast, LightSuite directly registered single-channel tdTomato fluorescence volumes from TRAP2×Ai14 mice without a dedicated autofluorescence channel. Despite sparse activity-dependent labeling^9^, background fluorescence provided sufficient anatomical contrast across tracts and boundaries to register brains accurately to the Allen CCFv3 and construct an average intensity template (Fig. 2G). Signal in each area reflected both Fos-dependent tdTomato expression and autofluorescence, the latter was visible in the anterior commissure and cerebellar white matter, with background intensity differing up to threefold across regions. Registering directly on reporter fluorescence, LightSuite thus removes the need for a secondary structural channel.

We benchmarked LightSuite’s registration against brainreg^17,18^, an established automated tool based on mutual-information registration. While brainreg achieved gross alignment on intact samples, it frequently misaligned local boundaries, cortical borders, and the hippocampus (Fig. 2H). To quantify registration precision against ground truth, we examined the primary somatosensory barrel cortex, whose distinct layer 4 cytoarchitecture is clearly resolved in autofluorescence (Fig. 2I, Ext. Data Fig. 5). LightSuite accurately positioned the barrel field within layer 4, whereas brainreg repeatedly misallocated barrels to incorrect cortical depths and produced significantly larger lateral displacement errors (Fig. 2J).

Evaluating whole-volume registration quality remains challenging. Traditional Dice overlap scores^19^ require labor-intensive manual segmentations, are restricted to coarse brain divisions, and do not scale across large cohorts. We therefore evaluated cross-animal registration consistency using an unsupervised metric: brains registered into a common coordinate space should show more strongly correlated regional fluorescence profiles across animal pairs. LightSuite yielded significantly higher pairwise correlations than brainreg, both globally and across substructures within every major brain division (Fig. 2K), confirming superior alignment consistency across cohorts.

### Fast cell detection across diverse areas

While deep-learning segmentation performs well on restricted fields of view, it is computationally costly across whole-brain volumes. LightSuite instead combines signal processing with lightweight machine learning. 3D band-pass filtering isolates cell-sized features while surrounding spatial frequencies estimate the local background (Fig. 3A). Dividing the band-pass volume by the local background converts raw intensity into a signal-to-background ratio (SBR), factoring out fluorescence variations across brain areas (Fig. 2G). Processing operates in GPU-accelerated 3D tiles, allowing full-volume execution on standard workstations without requiring global illumination flat-fielding^11^. To triage false positives, we trained a lightweight 2D convolutional neural network (CNN; Fig. 3B) that classifies candidates from orthogonal maximum-intensity projections (XY, XZ, YZ) of centered bounding boxes, bypassing the memory burden of 3D convolutions. The classifier achieved 98.7% validation accuracy, effectively eliminating bubble edges, tissue interfaces, and neurite varicosities (Fig. 3C). Across brains, the CNN discarded 10–20% of initial candidates (median 16%; Fig. 3D); rejected objects were systematically dimmer, smaller, and more elongated (Fig. 3E–G). The hybrid pipeline yields a highly specific soma population from only three 2D projections per candidate.

**Fig. 3.**
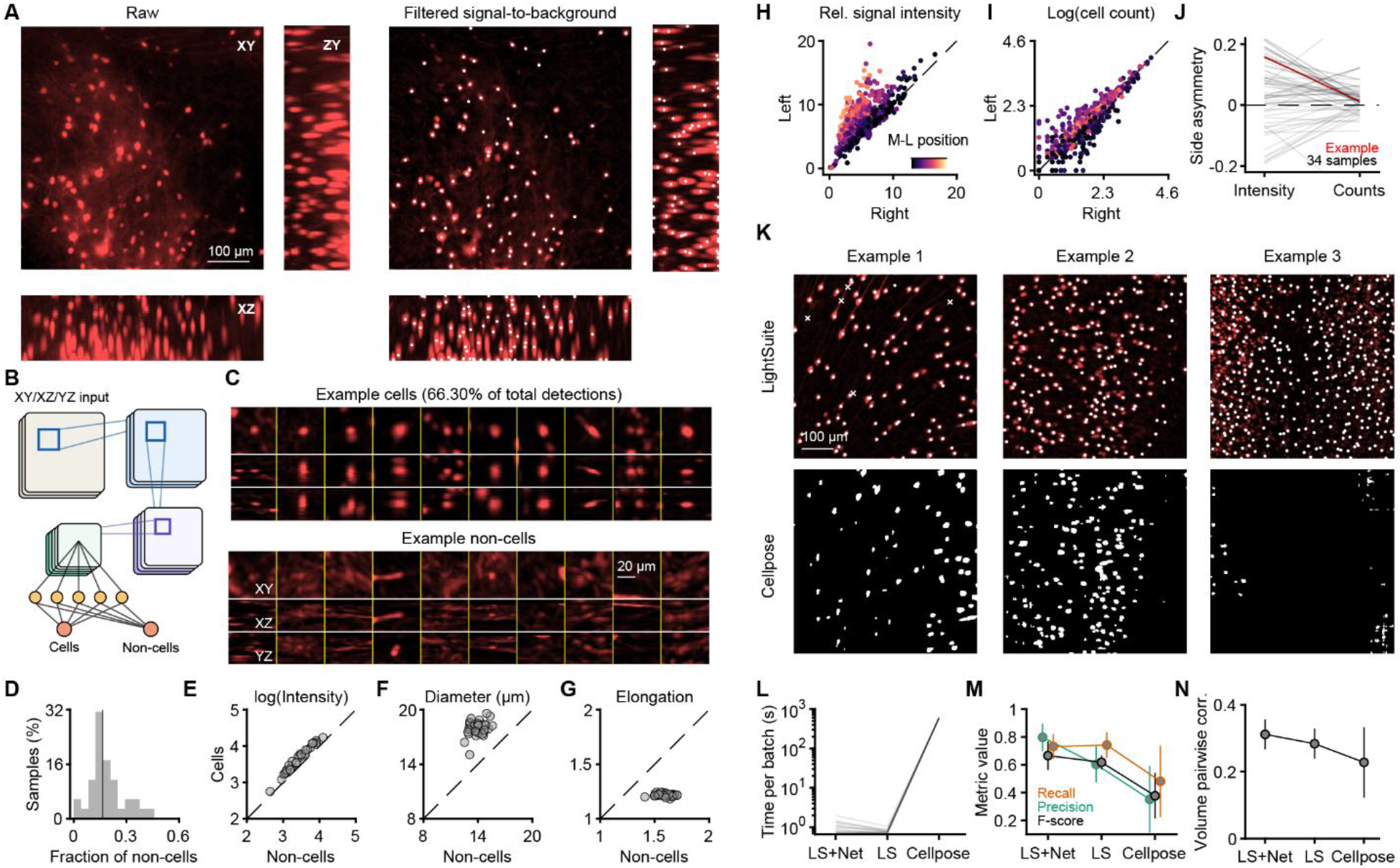
Fast cell detection across the entire mouse brain. **A**) Left: maximum-intensity projections of a tdTomato-labeled subvolume along three axes. Right: the same volume after band-pass filtering and division by the local background, which highlights cell-like objects. **B**) Schematic of convolutional network architecture for classifying image patches in three different views (maximum projections across XY, XZ, YZ) as cells. **C**) Ten example detections classified by the network as cells (top) and ten classified as non-cells (bottom), separated by yellow lines. For each cell and from top to bottom, XY, XZ and YZ projections are stacked vertically. The bar marks the scale of the XY projection. **D**) Distribution of the fraction of non-cells across multiple mouse brains (n=34). Black line marks the median at 0.16. **E**) Median intensity across detections classified as cells versus non-cells, for the same samples (logarithmic scale). **F**) Same as (E), but for the estimated diameter of the detection mask. **G**) Same as (E), but for elongation, defined as the ratio of the longest to the second-longest principal axis. **H**) Fluorescence intensity relative to background across Allen substructures (n=688), left versus right hemisphere, for an example sample illuminated with a unilateral light sheet; background is computed separately per hemisphere, and color encodes distance from the midline. **I**) As in (H), for cell counts (log scale) estimated by LightSuite within each area; paired comparison between left and right hemispheres shows no hemispheric bias (Wilcoxon signed-rank test, p=0.145). **J**) Side asymmetry index across samples (n=34), showing cell counts tightly centered around zero compared to intensities. Across samples, the absolute asymmetry index is significantly higher for intensities than for counts (Wilcoxon signed-rank test, p=1.1·10^-3^). **K**) Three example tissue subvolumes (maximum intensity projections) with different cell densities and backgrounds, illustrating detections for LightSuite (top) and segmentation results for Cellpose (bottom). Cellpose parameters, like those of LightSuite, were identical across batches. **L**) Processing time (logarithmic scale) per subvolume for LightSuite with (LS+Net) and without (LS) the classifier and for Cellpose. LS+Net was slower than LS alone (1.04 versus 0.77 s, Wilcoxon signed-rank test, p=1.6·10^-3^, n=16 subvolumes), and far faster than Cellpose (600.26 s, p=4.4·10^-4^). **M**) Benchmark metrics across manually annotated subvolumes (n=12). Recall (orange), precision (green), and F-score (black) for LS+Net, LS, and Cellpose. LS+Net achieves the highest precision (median 0.80) and F-score (median 0.67) while maintaining high recall (median 0.73). **N**) Pairwise cross-correlations between 3D intensity patches centered on detected somata across subvolumes (n=16). LS+Net yielded significantly higher patch correlation than LS alone (Wilcoxon signed-rank test, p=0.011) and Cellpose (p=8.4·10^-3^), reflecting higher morphological uniformity and fewer spurious detections. Error bars in (M) and (N) indicate 95% robust confidence intervals.

The one-sided light-sheet illumination we used here avoids the calibration of multi-sheet systems and the extra acquisition, calibration and fusion steps they require, but it imposes a cross-hemisphere asymmetry in raw fluorescence intensity (Fig. 3H) that grows with distance from the midline. LightSuite’s cell detection is resistant to this intensity variation: substructure cell counts are balanced across hemispheres (Fig. 3I) and far less asymmetrical than intensities (Fig. 3J). Counting cells rather than integrating intensity therefore removes an illumination artifact that would otherwise bias any hemispheric comparison.

We benchmarked LightSuite cell detection against Cellpose, a generalist approach that provides full cell segmentation^24^. No single Cellpose configuration worked across the brain: settings that segmented one area accurately often failed in another (Fig. 3K). LightSuite processed batches ∼600-fold faster than Cellpose (Fig. 3L). On manually annotated volumes, LightSuite recovered cell centers with recall and precision comparable to or higher than Cellpose (Fig. 3M). The median relative difference from manual counts was 7% (robust SD 30%) for LightSuite and 26% (robust SD 100%) for Cellpose. As an unsupervised quality metric, we evaluated pairwise cross-correlations among voxel neighborhoods around detected somata; genuine cells share stereotyped profiles, whereas false detections yield heterogeneous intensity correlations. LightSuite slightly outperformed Cellpose on this measure (Fig. 3N), and the classifier raised precision and overall performance over the base algorithm at a small cost in speed (Fig. 3L). LightSuite thus delivers segmentation-level precision at high throughput, identifying an average of 4.50·10^5^ (SD=1.19·10^5^) cells per mouse brain within 3–4 h on a desktop workstation.

### Consistent Fos activity mapping across individual mice

We next applied LightSuite to quantify brain-wide immediate-early gene expression across individual TRAP2×Ai14 mice, grouping cells at the substructure level of the Allen CCFv3. In these mice, tdTomato labeling was induced with 4-OHT following performance in visually guided tasks (Methods). Across substructures, tdTomato+ cell density correlated positively with standard anatomical neuron density from the Blue Brain Cell Atlas^38^ (ρ≈0.5; Fig. 4A,B), while absolute tdTomato+ densities were ∼100-fold lower than total neuron density. Across all samples, tdTomato+ density correlated more strongly with total neuronal density than with excitatory neuron density alone (Fig. 4B), consistent with Fos induction in both excitatory and inhibitory populations^39^. We used these density relationships as an independent biological benchmark: accurate atlas registration should align cellular distributions more faithfully with reference cytoarchitecture and yield higher inter-animal agreement. Brains registered with LightSuite showed significantly higher correlations to standard anatomical neuron density (Fig. 4C) and higher pairwise inter-animal correlation of substructure counts (Fig. 4D) than brains registered with brainreg, particularly across the isocortex and olfactory areas.

**Fig. 4.**
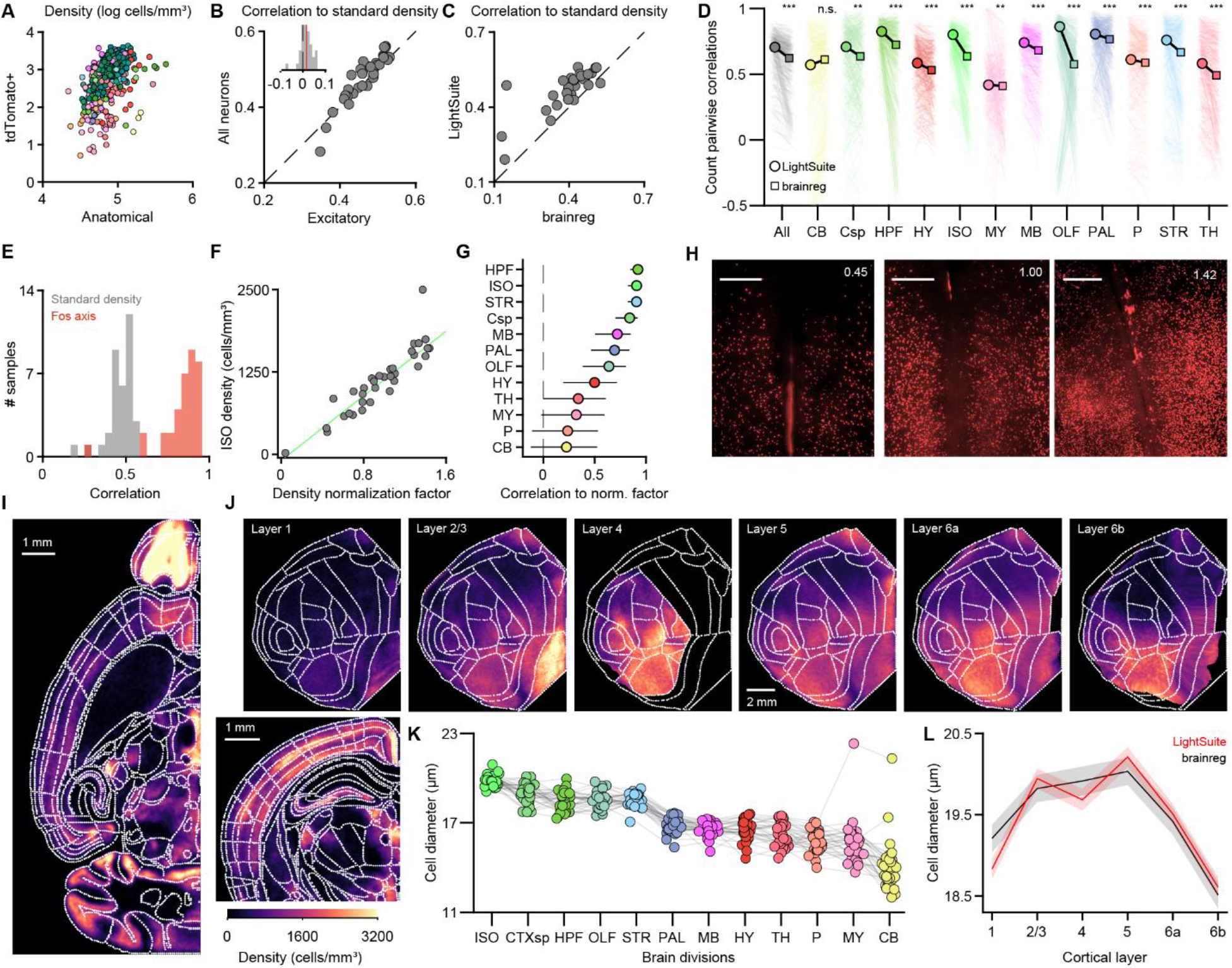
Substructure-resolved, cross-animal quantification of Fos-dependent tdTomato labeling with LightSuite. **A**) Standard anatomical density (logarithmic scale) of Allen Mouse Brain Atlas areas^38^ correlates with density of Fos-dependent tdTomato detections in the same area for an example mouse brain (n=429 substructures, Spearman’s ρ=0.53, p=1.3·10^-32^). **B**) The correlation between tdTomato+ density and the density of all neurons is higher than the correlation with excitatory neurons alone (n=34 brains, Wilcoxon signed-rank test, p=1.2·10^-3^). Inset shows the difference in correlation between the two groups and its median (red line). **C**) Correlation between tdTomato+ density and standard atlas density, for LightSuite versus brainreg (n=30 brains with brainreg registration, Wilcoxon signed-rank test, p=2.6·10^-6^). **D**) Pairwise correlations of counts in substructures of the Allen Mouse Brain Atlas, shown as in Fig. 2K. Across all areas (gray), LightSuite outperforms brainreg (n=435 mouse pairs, Wilcoxon signed-rank test, p<1·10^-50^). **E**) Distribution of Spearman correlation coefficients between tdTomato+ counts and neuronal density (gray) and between tdTomato+ counts and a common density profile extracted from all samples (orange). **F**) Projection strength onto the common density profile (density normalization factor) is approximately linearly related to tdTomato+ density in the isocortex (n=34, Pearson’s r=0.91, p=1.0·10^-13^). Green line is a robust linear fit. The point close to the origin corresponds to a brain with failed 4-OHT action. **G**) Pearson correlation coefficients between tdTomato cell density and the normalization factor across all brain divisions. Error bars are 95% confidence intervals. **H**) Maximum intensity projection axial view of three example samples in the same frontal area, each with a different estimated density normalization factor (numeric value in white). Scale bar is 500 μm. **I**) Top: example axial slice of the tdTomato+ cell-density map, averaged across animals after normalization (each animal weighted by its inverse density normalization factor). Bottom: a coronal cutout illustrating density across cortical layers. **J**) Cortical flatmaps of the tdTomato+ density map in (I), shown for each cortical layer. **K**) Estimated cell diameter across samples and brain divisions. Values have been multiplicatively normalized to the same median across samples to facilitate comparison across divisions. **L**) Estimated cell diameter across cortical layers comparing LightSuite (red) and brainreg (black) registration. Profiles and shaded 95% confidence intervals are estimated using a linear mixed-effects model across 43 cortical areas to account for area-specific baseline variation. The colormap of (I) and (J) is saturated to better illustrate density differences.

The tdTomato signal, however, reflects baseline and behaviorally relevant Fos expression on top of purely anatomical cell packing^40^. To isolate this baseline expression across animals, we extracted a single non-negative axis of expression by rank-one non-negative matrix factorization of the regional cell-density matrix across animals, independently of the anatomical reference^38^. This axis correlated substantially more strongly with each animal’s data (0.8–0.9) than anatomical neuronal density alone (Fig. 4E). Projecting each brain onto this axis yielded a per-animal normalization factor that scaled linearly with detected cell density in the isocortex and across divisions (Fig. 4F–H). This variation in labeling efficiency was not explained by animal weight (Ext. Data Fig. 6A); it may result from differences in effective 4-OHT delivery and should be factored out when pooling mice for a cohort average. Our data-driven normalization factor closely mirrored a standard-based factor derived from projecting sample densities onto the anatomical neuron density reference (Ext. Data Fig. 6B,C). However, scaling directly from anatomical neuron density left structured residuals across regions, reflecting baseline metabolic and behaviorally driven activation (Ext. Data Fig. 6D,E). These residuals peaked in the midbrain, hypothalamus, and isocortex (Ext. Data Fig. 6F). Within the isocortex, residuals revealed pronounced laminar specificity: layer 4 exhibited substantially higher Fos induction than predicted by cytoarchitecture, whereas layer 1 showed marked sparsity (Ext. Data Fig. 6G).

Normalizing each animal yielded an average brain-wide map of tdTomato+ cell density (Ext. Data Fig. 7), with prominent enrichment in the olfactory bulbs (Fig. 4I). Cortical flatmaps resolved distinct layer-specific patterns, with the highest density in visual cortex layer 4 (Fig. 4J). Layer 2/3 of the ventral retrosplenial cortex also showed high tdTomato+ density relative to its dorsal counterpart (Fig. 4J), which may reflect differences in cell-type composition across retrosplenial subregions^41,42^. Using the segmentation accompanying each cell detection, we also estimated soma diameter; it varied consistently across major divisions, was highest in the isocortex, cortical subplate and hippocampus and lowest in the hypothalamus and cerebellum (Fig. 4K). Within the isocortex, soma diameter peaked in layers 5 and 2/3 (Fig. 4L). In absolute terms, our estimated diameters (∼20 µm) were moderately higher than true anatomical soma sizes, typically between 10 and 16 µm^38,43^. That overestimate may arise from the SBR-based dilation of candidate regions and from point-spread-function broadening in undeconvolved light-sheet acquisitions. Nevertheless, the relative laminar profiles faithfully reproduce canonical cortical cytoarchitecture, where pyramidal somata in layers 5 and 2/3 are systematically larger than those in layers 4 and 6^43–46^. Registration with brainreg, by contrast, obscured this laminar size hierarchy because cortical layers were misallocated. LightSuite thus extracts biologically structured cellular metrics that extend beyond simple regional counts.

### Spinal cord registration

Despite its central role in sensorimotor coordination^47^, the spinal cord is rarely integrated into automated CNS pipelines because cleared cords undergo severe, non-uniform bending and torsion^48^. To map spinal volumes into standardized coordinates, we extended LightSuite’s landmark framework to cord morphology. We cleared and imaged spinal cords from mice expressing membrane-anchored EGFP under the CNP promoter (CNP-mEGFP), which provides endogenous contrast between white matter and gray matter without antibody staining (Fig. 5A). Because tissue deformation varies continuously along the rostrocaudal axis, LightSuite estimates centerline curvature and axial torsion by fitting smoothing splines to user annotations across sparsely sampled axial slices (Fig. 5B). These trajectories computationally straighten and untwist the volume into a canonical cylinder (Fig. 5C). A multi-metric non-rigid fit incorporating user-defined control points then aligns the straightened specimen to a reference spinal cord atlas^49^ (Fig. 5D).

**Fig. 5.**
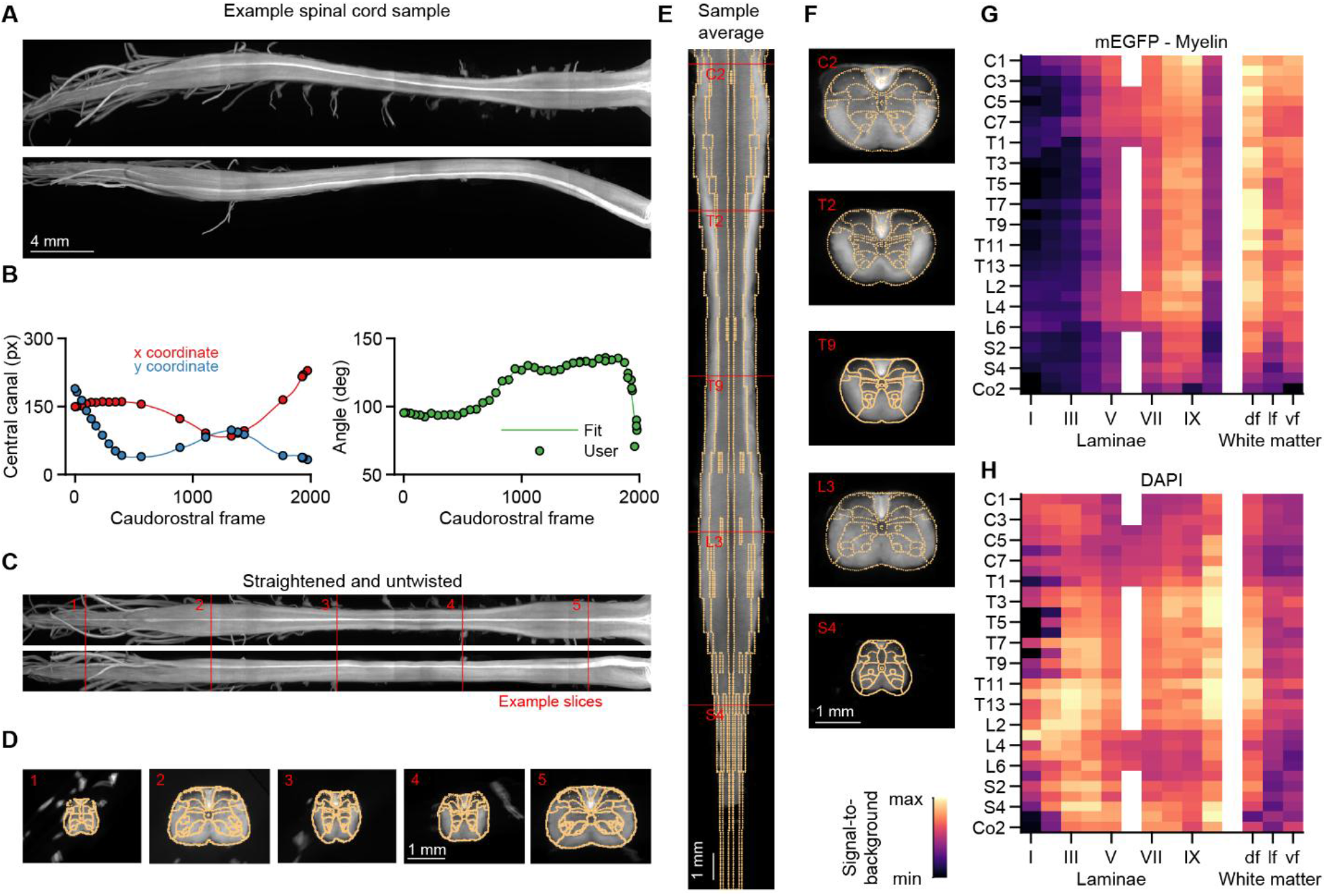
Untwisting and atlas registration of cleared mouse spinal cords. **A)** Maximum-intensity projections (top: x/z, bottom: y/z) of a cleared spinal cord labeled with CNP-mEGFP, which captures mature myelinating cells. **B**) User-annotated coordinates for the central canal (left) and dorsoventral axis (right) fitted with penalized smoothing splines. **C**) Computationally straightened and untwisted volume (top: x/z; bottom: y/z); red lines mark the positions of the cross-sections shown in (D). **D**) Axial cross-sections of the straightened sample at those positions, with the atlas fit overlaid (yellow). **E**) Average myeloarchitectural map across five spinal cords (red lines mark slices in (F)). **F**) Representative cross-sections from cervical (C), thoracic (T), lumbar (L), and sacral (S) segments. **G**) Myelin (mEGFP) signal-to-background across spinal segments and anatomical subdivisions. **H**) As in (G), for nuclear DAPI staining. df, dorsal funiculus; lf, lateral funiculus; vf, ventral funiculus.

We validated registration accuracy by constructing an average myeloarchitectural map across five spinal cords (Fig. 5E,F), which exhibited sharp intensity boundaries aligned with atlas annotations. Segmental quantification across laminae and funiculi^50^ identified the highest myelin density in the dorsal funiculus and revealed a dorsal-to-ventral laminar gradient, increasing from lamina I to lamina IX (Fig. 5G). Nuclear DAPI counterstaining instead showed enriched cellularity in gray matter relative to white matter across all segments (Fig. 5H). Applying LightSuite to a cleared spinal cord from a PDGFRα-CreER[T2]^tg/+^ mouse revealed higher oligodendrocyte-precursor-cell-derived tdTomato+ cell density in white matter funiculi than in gray matter laminae, consistent with known white matter precursor cell enrichment^51^ and elevated proliferation^52^. LightSuite also detected tdTomato+ cells outside the cord boundaries within peripheral tissue protrusions corresponding to spinal nerve roots. LightSuite thus computationally rescues distorted cord specimens and achieves registration precision sufficient for substructure-level quantification of intensities and cell densities.

### Registration and cell detection in brain slices

Because much neuroscience data relies on conventional wide-field microscopy of physical sections, we built a slice-processing module into LightSuite. Existing section-based tools^20,53–55^ rarely assemble true 3D volumes for cohort averaging or allow users to arbitrarily edit nonlinear transforms without re-optimizing; they also can fail on heavily damaged slices^30^. LightSuite lets users systematically order, rigidly align, and nonlinearly fit 2D sections into a 3D volume in atlas coordinates (Fig. 6A), filling gaps from irregular spacing by nearest-neighbor interpolation. Compared to 3D imaging, the slice workflow is more involved because each section carries its own transforms, but LightSuite applies the same active-learning principle to find the best-matching atlas slice on which a user can annotate landmarks.

**Fig. 6.**
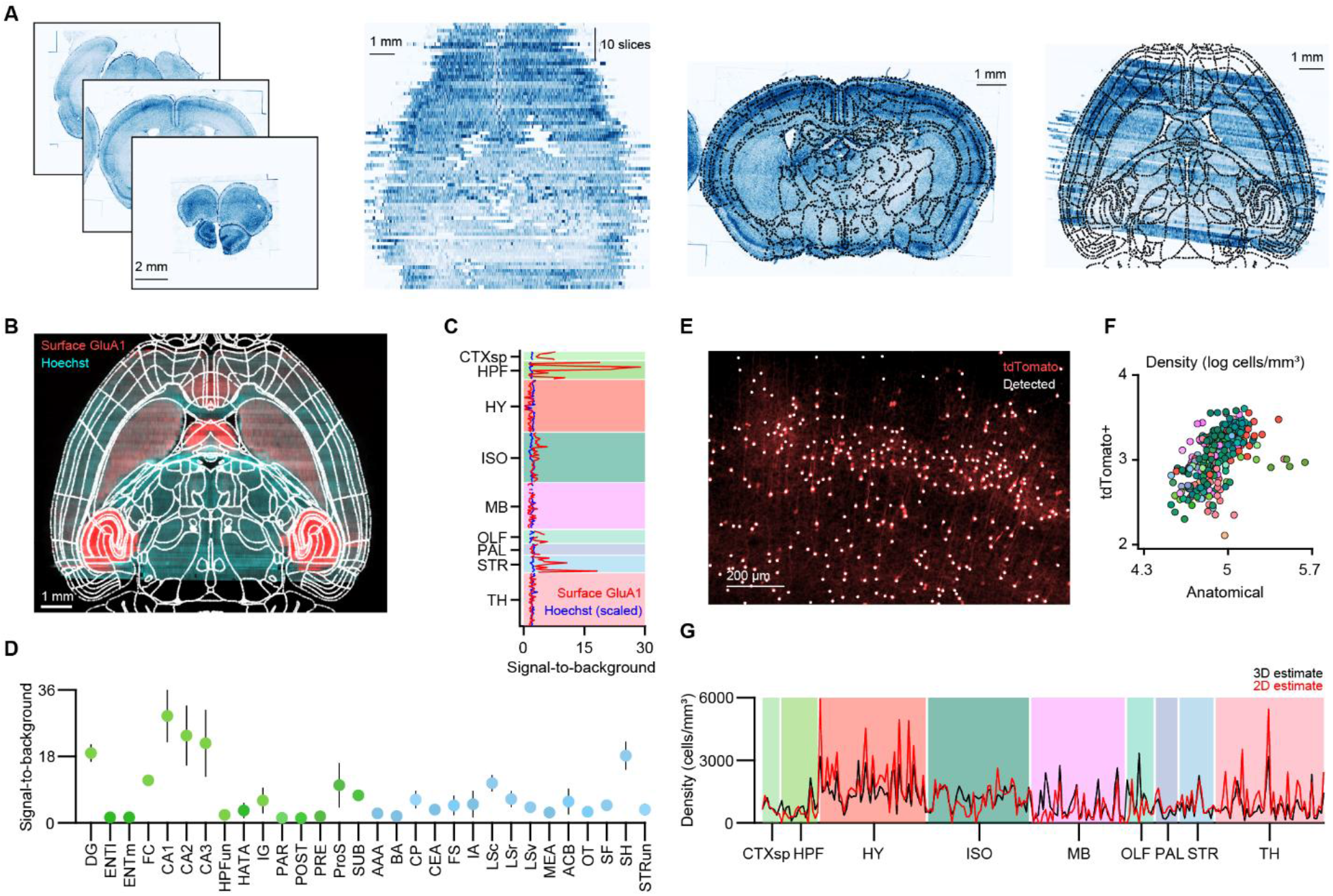
Volumetric reconstruction and quantification from serial histological sections. **A**) Reconstruction workflow for coronal wide-field sections (left; DAPI). Slices are ordered along the rostrocaudal axis, rigidly aligned at equidistant intervals (middle-left), interactively adjusted for cutting angle, and non-rigidly registered using landmark pairs (middle-right). Transforms are inverted to assemble a 3D volume in atlas space (right). **B**) Average 3D reconstruction of five brains stained for Hoechst and surface GluA1 (nanobody-stained SEP-GluA1). **C**) Median intensity profiles (n=5 mice) across brain areas. To ease visual comparison, Hoechst intensity is multiplicatively scaled to match the median intensity of surface GluA1 across areas. **D**) Median intensity signal profile (n=5 samples) for individual areas grouped by brain division, showing that surface GluA1 is strongly enriched in the hippocampus (green shades) and striatum (cyan shades). Error bars are robust standard deviations. **E**) Detected tdTomato+ cells (white dots) on a coronal slice cutout. **F**) Regional tdTomato+ cell density from wide-field sections correlates with reference anatomical neuron density (n=312 substructures, Spearman’s ρ=0.62, p=3.1·10^-34^). Both are plotted in logarithmic scale. **G**) Median tdTomato+ cell densities from serially sectioned brains (n=3) predict the light-sheet 3D estimate (n=33) across regions (n=235 substructures, R^2^=0.48).

Applying the slice module to coronal sections stained for the surface AMPA receptor subunit GluA1 (Methods), we reconstructed brain-wide molecular density volumes (Fig. 6B). These maps confirmed marked GluA1 enrichment in the hippocampus and striatum, and identified prominent localization in the lateral septum (Fig. 6C,D). Extending SBR detection to 2D sections (Fig. 6E) yielded regional tdTomato+ cell densities that strongly correlated with standard anatomical neuron densities (Fig. 6F). Moreover, regional cell densities estimated from 80-μm physical sections predicted whole-brain 3D light-sheet measurements (Fig. 6G). The slice module thus recovers quantitatively consistent molecular and cellular maps from standard histology, bridging physical sectioning and 3D atlas frameworks.

### Registration of functional ultrasound volumes

LightSuite’s registration framework is not limited to fixed tissue. *In vivo* fUS has emerged as a powerful tool for brain-wide hemodynamic mapping, but cohort-scale studies require reliable alignment of repeated recordings across sessions and animals into a common reference space. Alignment is challenging because fUS contrast reflects vascular structure rather than cytoarchitecture, spatial resolution is coarse (typically ≥100 μm), and cranial windows introduce non-uniform soft-tissue deformations (Fig. 7A). To date, volumetric fUS registration has largely remained manual, relying on affine transformations adjusted by stretching and shearing individual axes to visually match anatomical boundaries^21,56^. While different sessions from the same animal can be automatically realigned, registering across animals and to a reference vascular atlas remains challenging, requires a trained eye, and is a time-consuming step. Automated vascular registration approaches have been proposed^57,58^, but they have not been validated against *in vivo* functional activation maps or tested on low-resolution volumetric array fUS, and not all proposed approaches are open-source. LightSuite bridges this appearance gap across both planar motorized scans^21^ and volumetric matrix arrays^59^. We first registered a high-resolution power-Doppler vascular template^21^ into the CCFv3 to transfer atlas annotations onto vascular contrast. Within each mouse, functional sessions were rigidly aligned to a single recording designated as its anatomical reference, which was then registered to the vascular template via affine transformation followed by landmark-guided non-rigid B-spline deformation (Fig. 7B). To prevent overfitting to coarse vascular fluctuations, we constrained the B-spline grid to 1.6-mm isotropic spacing and guided alignment with a median of 219 user landmarks per mouse.

**Fig. 7.**
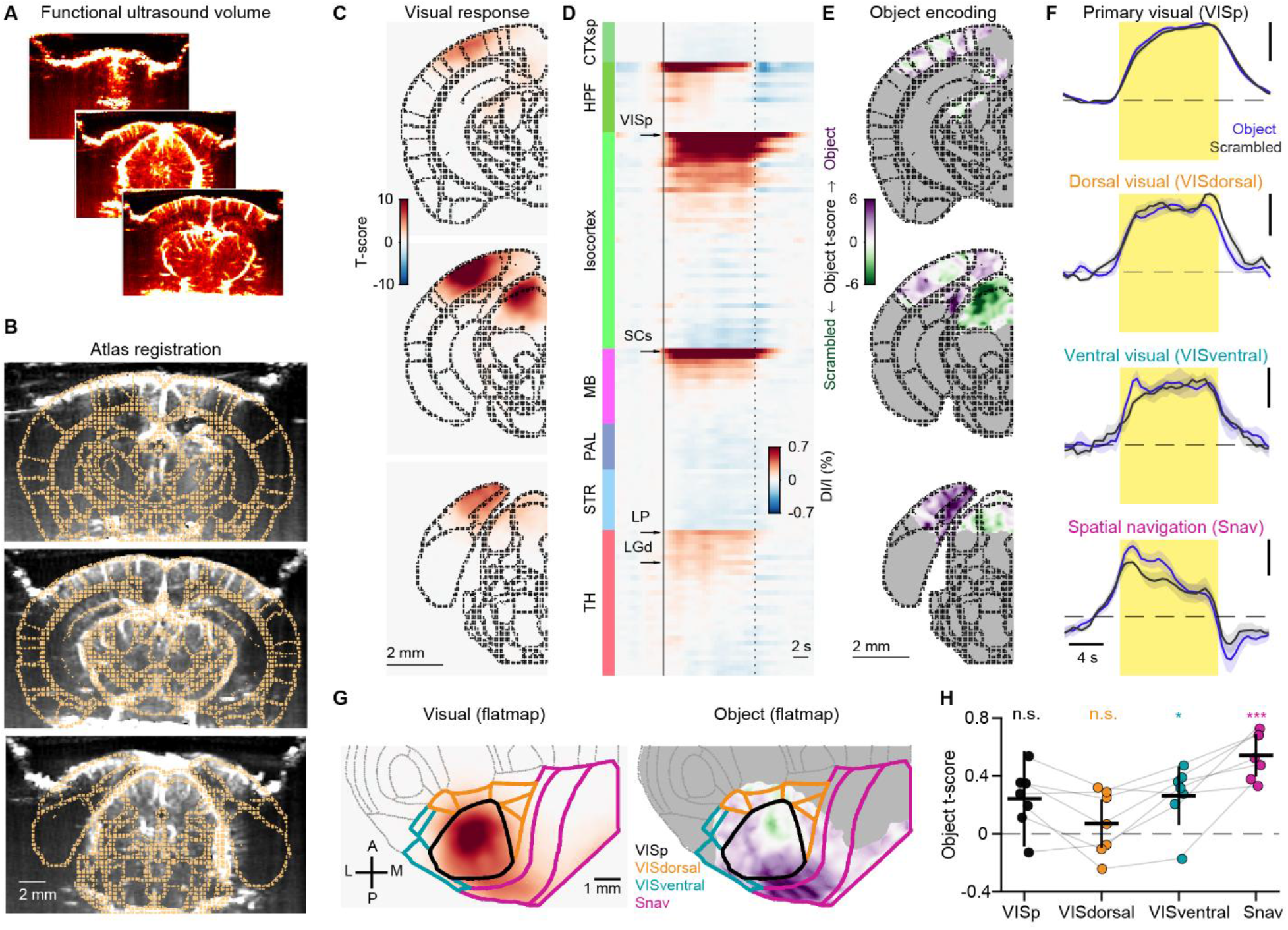
Application to functional ultrasound (fUS) imaging. **A**) Illustration of three coronal slices from a fUS volume. **B**) Non-rigid fit of the Allen Mouse Brain atlas (yellow dots) to an anatomical fUS scan, here shown across three coronal slices using LightSuite. Note that the atlas boundaries are shown at the structure level. **C**) Average visual t-score maps (Methods) in three representative coronal slices, across seven mice registered to the atlas. Voxels outside the atlas were set to zero. **D**) Stimulus-triggered traces of the relative change in signal intensity (ΔI/I), averaged across the voxels of each atlas structure. Structures smaller than 0.1 mm^3^, and those of the hypothalamus, pons and cerebellum, are omitted. Within each brain division, structures are sorted by their mean response during the stimulus window. Arrows mark four structures of the visual pathway (VISp, SCs, LP, LGd). **E**) Group object t-score map (object versus scrambled contrast; Methods) in the same three coronal slices as in (C). Purple, preference for objects; green, preference for scrambled images; gray, visually unresponsive voxels. **F**) Normalized object (blue) and scrambled (black) ΔI/I traces for four candidate area groups: primary visual, dorsal higher visual, ventral higher visual and spatial navigation. Before averaging, each structure was normalized to its peak response value to the combined visual stimulus. Yellow shaded regions mark the stimulus window; the vertical and horizontal scale bars denote normalized ΔI/I and time, respectively. **G**) Cortical flatmap projections (posterior cortex) of the visual (left) and object-encoding (right) maps, with the same color scales as in (C) and (E). The four area groups are outlined in color; gray, visually unresponsive cortex. **H**) Object t-score per area group (n = 7 mice). For each mouse, the t-score was summarized as the median over the visually responsive voxels of each atlas area (16 areas with ≥10 responsive voxels) and averaged over the areas of each group. Dots show individual mice, gray lines connect the same mouse across groups, and black bars show the population mean ± 95% CI from a linear mixed-effects model (Methods). Group effect (likelihood-ratio test): χ²(3)=15.4, p=0.002. Comparison to zero (Wald test, FDR-corrected across the four groups): VISp p=0.196, VISdorsal p=0.390, VISventral p=0.022, Snav p=1.8·10^-10^. Pairwise comparisons (Wilcoxon signed-rank test): Snav vs VISdorsal p=0.016, VISdorsal vs VISventral p=0.032. *p < 0.05, ***p < 0.001; n.s., not significant.

We evaluated LightSuite on *in vivo* volumetric array fUS recordings from mice presented with visual object and scrambled control images^59^. In atlas space, regional hemodynamic traces faithfully captured visual activations across the early visual pathway, including primary visual cortex (VISp), the dorsal lateral geniculate and lateral posterior thalamic nuclei, and the superior colliculus (Fig. 7C,D). To rigorously benchmark non-rigid deformation, we compared it not to crude manual stretching, but to an affine transform optimized directly within LightSuite using joint image contrast and the curated user landmarks. Even against this optimized affine baseline, non-rigid registration significantly improved alignment across every quantitative metric: it suppressed visual activation leakage across atlas borders, increased bilateral hemispheric response correlations, and boosted pairwise inter-animal correlations for both vascular anatomy and visual activation maps (Ext. Data Fig. 9A–F). Furthermore, non-rigid registration unmasked stronger object-related recruitment in the postsubiculum (Ext. Data Fig. 9G,H) and resolved fine spatial organization in cortical object sensitivity (Fig. 7E–H) that was obscured under manual affine registration^59^. Object sensitivity was higher in ventral than in dorsal visual areas^60,61^, and the cortical flatmaps revealed a spatial gradient of object preference within the VISp (Fig.7G). LightSuite thus extends unified atlas mapping to *in vivo* functional hemodynamics, enabling reproducible cohort analysis across sessions and animals.

## DISCUSSION

LightSuite removes the computational bottlenecks that hinder cohort-scale mapping of the CNS, and with it we found that the efficiency of Fos-dependent labeling varies several-fold between animals along a single axis that is separable from anatomy. By coupling damage-resilient point-cloud initialization with user-steered landmark optimization, LightSuite registers mouse brains to the Allen CCFv3 more consistently than an established automated pipeline. Its signal-to-background cell detection is more than two orders of magnitude faster than a generalist deep-learning approach^24^, and it enables robust cellular quantification on a standard workstation. The same framework untwists cleared spinal cords, assembles serial 2D histological sections into 3D volumes, maps fiber trajectories, and non-rigidly registers *in vivo* fUS volumes. Applied to previously published recordings^59^, LightSuite also resolved a ventral-versus-dorsal dichotomy in mouse visual object sensitivity^60,62^. LightSuite thus bridges cellular-resolution histological snapshots with mesoscale hemodynamics.

Registration quality across cohorts requires evaluation standards that move beyond manual Dice overlap scores^19,31,63,64^. To provide an objective anchor, we introduced a ground-truth metric based on the displacement of the stereotyped layer 4 barrel field of the primary somatosensory barrel cortex. In parallel, we established unsupervised cohort-scale benchmarks: cross-animal correlation of regional tdTomato fluorescence, and agreement between measured cell densities and standardized anatomical references^38^. These metrics convert registration evaluation from expert outlining into an automated, cohort-scalable benchmark of alignment fidelity. Neither the LightSuite’s registration nor the unsupervised cross-animal correlation benchmark is specific to the brain. Both require only a reference template, so in principle they apply to any organ for which such a template exists^4^, and the spinal cord registration already extends the approach beyond the brain.

Four limitations currently bound LightSuite. First, registration accuracy depends on user landmarks, which makes it operator-dependent; inter-annotator variability remains to be quantified. The active-learning interface lowers the burden, and the landmark residual provides a per-sample quality figure; a promising remedy is a learned keypoint detector^65^ that would make placement reproducible and remove the manual step. Second, LightSuite detects rather than segments cells. Because estimated neuronal diameters (∼20 µm) exceed anatomical soma sizes, our findings are restricted to relative cross-region and layer comparisons, with densely labeled regions risking cell merging; upstream deconvolution or a dedicated segmentation model would address both issues. Third, photon attenuation in deep and ventral structures stems from our acquisition setup and is amplified by the unilateral light-sheet illumination we adopted to keep cohort-scale storage tractable. Deep subcortical tdTomato+ counts must be interpreted cautiously, though dual-sided illumination could resolve this issue at the cost of scan time. The brain-wide Fos expression atlas we provide as a general reference (Ext. Data Fig. 7) inherits the same ventral attenuation, and it also reflects labeling induced under photopic conditions, which drive immediate-early gene expression in the visual cortex^9,66^; induction in darkness would provide a complementary reference. Finally, spinal cord and fUS analyses are limited by available templates: the spinal cord atlas^49^ lacks the granularity of brain atlases, and the vascular template^21^ paired with ∼150 μm fUS voxels limits interpretable structure size. Both constraints will relax as higher-resolution templates appear and spatial resolution improves, and LightSuite accepts any template supplied in the same format, and thereby maintains a major advantage in the face of evolving datasets and atlases.

## METHODS

### Experimental animals

For activity-dependent labeling experiments, we used 37 Fos2A-iCreER×Ai14(RCL-tdT)-D mice (four females), aged 8 to 35 weeks, with the crossing done between TRAP2^9^ (JAX:030323) and Ai14^67^ mice (JAX: 007914). For surface AMPA receptor subunit GluA1 labeling, we used five mice (two females) from the Gria1^tm8.1Rlh^/J SEP-GluA1 KI mouse line^68^(JAX:037254). We additionally used nine male wild-type (C57BL/6J) mice that had been implanted bilaterally with optic fibers. All experiments were carried out in accordance with the Institutional Animal Care and Use Committee of the University of Geneva and in compliance with the Swiss Veterinary Law Guidelines, approved by the ethics committee of the Cantonal Veterinary Office of Geneva (GE191, GE519). Mice were housed in groups of up to five per cage under a 12/12-h non-inverted light/dark cycle with ad libitum access to food and water. The ambient temperature in the animal facility was 23 °C and the relative humidity was maintained around 50%.

For the registration and analysis of spinal cord samples, five male mice aged 8–14 weeks were used. All mice carried the CNP-mEGFP transgene (JAX:026105), which enables fluorescent labeling of mature oligodendrocytes and myelin. Four out of five mice also carried additional transgenes. The exact genotypes are as follows: CNP-mEGFP^tg/tg^ (one mouse), PDGFRαcreERT^tg/+^;Ai14^KI/+^;CNP-mEGFP^tg/tg^ (one mouse) and Tmem117^fl/+^;CNP-mEGFP^tg/tg^ (three mice). All animal care and experimental procedures were conducted in accordance with the Swiss National Institutional Guidelines of Animal Experimentation (OExA; 455.163) and under license approval ZH35166, issued by the Kanton Zürich Gesundheitsdirektion Veterinäramt (Zurich, Switzerland). Mice were housed in groups of up to five animals per cage in a temperature-controlled room under a 12 h light/dark cycle, with ad libitum access to water and standard laboratory chow (Kliba, Nafag).

### Animal behavior, surgery and pharmacology

Behavioral history varied across the cohort: two mice performed a head-fixed foraging task, four mice remained in their home cage, and the remaining mice performed a freely moving visual decision-making task based on two-alternative location discrimination. Half of the latter mice performed the task jointly with a conspecific. Task-trained mice were water-restricted and extensively trained before induction. In every case, 4-OHT was injected under photopic conditions and only after the behavioral session; mice were then placed individually in a dark box for at least one hour to limit additional sensory input during the labeling window. The nine wild-type mice used to demonstrate fiber-track reconstruction (Ext. Data Fig. 1) were prepared for optogenetic inhibition: each received bilateral intracranial injections of AAV-hSyn-stGtACR2-FusionRed, followed by bilateral implantation of 400-μm-diameter optic fibers over the injection sites.

To induce Cre-recombinase expression in TRAP2 mice, we injected a lipid-based 4-OHT solution i.p. as previously described^9^. Briefly, 4-OHT (Sigma-Aldrich) was dissolved in ethanol (20 mg/mL) for long-term storage at −20 °C. On the day of induction, a 150-μL aliquot was thawed, sonicated to promote suspension, and mixed with 200 µL sunflower seed oil (Sigma-Aldrich) before re-sonication. The mixture was incubated at 37 °C on a shaker to evaporate residual ethanol, yielding a 4-OHT concentration of 15 mg/mL. Mice were briefly anesthetized with isoflurane immediately before injection.

### Histology, brain clearing and immunohistochemistry

Mice were anesthetized with 3–4% isoflurane and euthanized with an i.p. injection of pentobarbital (Esconarkon, 150 mg/kg), preceded by an injection of buprenorphine (0.1 mg/kg) for analgesia. They were then transcardially perfused using a peristaltic pump with 20–30 mL of 0.01 M PBS, pH 7.4, and then with 20–30 mL of 4% paraformaldehyde (PFA) in phosphate buffer (PB; pH 7.4). The brains were post-fixed at 4 °C in the same PFA solution for 12 to 48 h and then transferred into PBS.

For optical clearing of whole mouse brains, we used the EZ Clear protocol^69^. Fixed brains were delipidated in 50% tetrahydrofuran for 36 to 48 h, washed thoroughly in distilled water, and incubated in refractive-index-matching solution (EZView) for at least 72 h. Spinal cord samples were cleared using SHIELD with DAPI counterstaining^70^. The tissue was then immobilized in an agarose gel and stored in 100% EasyIndex solution until imaging.

For slice experiments, after PBS washing, brains were embedded in a 4% agar-agarose gel before cutting slices at 80 or 150 µm thickness with a vibratome (VT 1000 S, Leica). SEP-GluA1 mouse sections were incubated with slight agitation (40 rotations/min) for 2 h at room temperature in a saturation solution containing a mix of 5% BSA and NGS, and PBS. Brain sections were then incubated with slight agitation (40 rotations/min) overnight at 4°C with an Alexa Fluor 647-conjugated alpaca anti-GFP VHH nanobody (ChromoTek GFP-Booster Alexa Fluor® 647, Proteintech; 1:500–1:1000) in the same blocking solution. Sections were washed three times for 15 min in PBS before a Hoechst solution (dilution 1:1000, Invitrogen, #33342) was used for fluorescent nuclear counterstaining. Slices were incubated for 10 min in that solution with slight agitation then washed twice for 5 min in PBS. If fluorescent nuclear counterstaining was not carried out using Hoechst solution, a mounting medium containing DAPI (Vectorlabs, H-1500) was used.

### Light-sheet microscopy

Cellular-resolution light-sheet imaging was performed using a customized version of the Clarity Optimized Lightsheet Microscope^71^. The samples were illuminated tile-by-tile by a single digitally scanned light-sheet, using a 488- or a 561-nm wavelength laser. Two excitation objectives (4×, NA 0.28) were used to form the Gaussian-shaped light-sheet. A self-adaptive positioning system maintained image quality over up to 1 cm of tissue during z-stack acquisition, compensating for refractive-index inhomogeneity along the stack. Illumination and detection were synchronized by locking the scanning beam to the unidirectional readout of an sCMOS camera chip, producing a virtual-slit effect that improves image quality. Emitted fluorescence was filtered (525/50 nm, 609/54 nm and 680/42 nm Semrock BrightLine HC) and imaged on an ORCA-Flash 4.0 V3 CAMERALINK digital CMOS camera at 4 fps as 16-bit images (2048×2048 pixels), in rolling shutter mode. To obtain whole-brain volumes, 3D tiles were stitched with TeraStitcher^72^.

Spinal cord samples were imaged using a mesoSPIM (mesoscale selective plane illumination microscopy) V6 light-sheet microscope^8^. The samples were mounted in the immersion chamber with the dorsal and ventral sides oriented toward the illumination lasers. For image acquisition, simultaneous dual-sided illumination was used in combination with a Mitutoyo M Plan Apo 2×/0.055 objective. The spinal cord was imaged as a grid of 1×5 tiles, resulting in a pixel size of 2.75 × 2.75 µm in the x- and y-dimensions and 5 µm in the z-dimension. Illumination was performed using 488-nm lasers at 50% power and 405-nm lasers at 80% power in combination with an exposure time of 20ms. Fluorescence emission was detected using a Quadband 405/488/561/640 filter and Hamamatsu ORCA Lightning sCMOS camera. Spinal cord volumes were stitched with BigStitcher^73^.

Optic-fiber-implanted mouse brains were also imaged with a mesoSPIM light-sheet microscope, but illumination was delivered unilaterally. Two-channel volumetric imaging was performed sequentially using 488-nm and 561-nm excitation laser lines. Emitted fluorescence was collected through a 530/43-nm bandpass filter for the 488-nm channel and an LP561 long-pass filter for the 561-nm channel. Samples were acquired at 1× or 1.25× zoom (in-plane pixel size: 6.55 or 5.26 μm) with an axial step size of 5 μm.

### Wide-field microscopy

Slices were mounted on Superfrost microscope slides (Epredia) with mounting medium (Fluoromount) and covered with 24 × 60 mm coverslips (Menzel-Gläser). All photomicrographs from slides with brain slices were taken with a Zeiss Axio Scan.Z1 with 10× objective at the bioimaging platform of the University of Geneva. Individual photomicrographs from the same sample were combined into an overview image by stitching algorithms with shading correction. Stitching was performed in the Zeiss software (10% tile overlap, 5% minimal overlap and 10% maximum shift) using the nuclear staining channel. In a subset of brains, slices were not ordered at imaging acquisition. Using the annotation GUI of LightSuite’s slice module, we ordered coronal slices along the anteroposterior axis and flipped them to a consistent mediolateral direction. Slices for which stitching failed, or that showed excessive staining artefacts or folding, were excluded from further processing using the same annotation GUI.

### Volumetric data preprocessing

For each imaging plane, a representative background intensity was estimated as the mode of the non-zero values within a random subsample of up to 20000 pixels drawn from that plane, and all zero-valued (empty or padding) pixels were set to this value so that background regions took a representative intensity rather than a hard zero. Each background-corrected plane was then downsampled in x–y by bicubic interpolation and accumulated into a stack; once all planes had been processed, this stack was downsampled along z by linear interpolation to yield the registration volume at 20 µm isotropic voxel spacing. This procedure was applied to every channel, and the resulting per-channel registration volumes were saved as TIFFs. For channels designated for cell detection, each background-corrected, full-resolution plane was additionally median-filtered (3×3 kernel, symmetric padding) to suppress acquisition artifacts.

### Volumetric cell detection

Cell detection was performed on the full-resolution detection channel in overlapping 3D batches sized to fit GPU memory (1800x1800x32 voxels), with adjacent batches overlapping by a 4*r_cell_* margin to avoid edge effects. Here 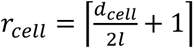 is an approximate cell radius in voxels derived from the user-specified cell diameter *d_cell_* and the minimum side of a voxel *l*. For all of our brain samples we set *d_cell_* to 14 μm and *l* was 1.44 μm.

#### Bandpass filtering

Each batch was transferred to the GPU in single precision and bandpass-filtered in 3D with a 4th-order Butterworth filter applied in the Fourier domain. The filter was defined around a corner frequency *f_c_* = 2⁄(*πr_cell_*) cycles per pixel, with the high-pass cutoff set to *f_c_*/2 and the low-pass cutoff to 5*f_c_*. To keep the cutoffs invariant to padding, frequency-space distances were normalized by the unpadded batch dimensions while the FFT itself was computed on a grid zero-padded to the next power of two along each axis; the batch was symmetrically padded before transformation to suppress wraparound artifacts and unpadded afterward. Frequency distances were scaled per axis by the voxel anisotropy ratio so that the filter response is isotropic in physical space.

#### Bandpass-normalized detection

A background estimate was formed by subtracting the bandpass-filtered batch from the original batch. To guard against saturated or empty regions, this background was clipped between the 10th and 90th percentiles of its positive values (estimated from a random subsample of up to 30000 voxels) and then smoothed with a 3D Gaussian (σ = 2 × the per-axis anisotropy ratio). Dividing the bandpass-filtered batch element-wise by this background produced a signal-to-background-ratio (SBR) volume. Candidate cell centers were voxels that simultaneously were local maxima of the SBR volume within a window of 4σ per axis, exceeded the primary SBR threshold (default 0.5), and had a bandpass amplitude above the 98th percentile of the sampled bandpass values. Each accepted maximum was then dilated with a cuboid structuring element (side length = ⌈4 · σ · anisotropy⌉ + 1 voxels per axis), and all voxels connected to a peak whose SBR exceeded the secondary threshold (default 0.3) were aggregated into a candidate region using 18-connectivity. The primary and secondary SBR thresholds and the amplitude percentile were set heuristically on pilot volumes, to retain dim somata while keeping the number of candidates passed to the classifier tractable. Lower thresholds increase recall at the cost of many more candidates for the CNN to evaluate; all three are user-adjustable parameters.

#### Morphological filtering

Connected components of candidate voxels were characterized with MATLAB’s regionprops3 to obtain the weighted centroid, principal-axis lengths, equivalent diameter, and mean intensity of each region. A region was rejected if the ratio of its first to second principal-axis length exceeded 2.5 (elongated structures such as neurite fibers), or if its anisotropy-corrected equivalent diameter was smaller than *r_cell_* (sub-cellular fragments) or larger than 8*r_cell_* (large bright artifacts such as tissue folds). To prevent double-counting across batches, detections whose centroids fell within the batch overlap margins were discarded. Surviving centroids were mapped back to global volume coordinates, their equivalent diameter was converted to physical units by multiplying by the cube root of the voxel volume, and each detection was stored together with its mean intensity, physical equivalent diameter, and principal-axis elongation ratio.

### Cell classification

A proportion of surviving detections are false positives arising from bubble edges, tissue–solution interfaces, or bright neurite varicosities. To triage these artifacts, we trained a lightweight convolutional neural network (CNN) in MATLAB. Rather than classifying 3D volumes, which would substantially increase storage and computational requirements for the training set, the network operates on maximum-intensity projections along each of the three spatial axes, centered on the candidate centroid (patch size = 4*r_cell_*). The three 2D projections are stacked as channels of a single input image and processed by a CNN comprising three convolutional blocks with 16, 32, and 64 filters respectively (3×3 kernels, same padding), each followed by batch normalization and a Rectified Linear Unit (ReLU) activation; the first two blocks include 2×2 max-pooling with stride 2. The resulting feature maps feed a fully connected layer with two outputs and a softmax function, trained end-to-end with cross-entropy loss. Training data were balanced between cells and non-cells, and augmented online with 3D-consistent random reflections: each spatial axis was independently flipped with 50% probability, with the corresponding image axes of all three views transformed jointly to preserve geometric consistency. The network was trained on 6872 manually labeled examples with an 80/20 training/validation split, using the Adam optimizer for 100 epochs with a mini-batch size of 64 and shuffling at every epoch. It achieved 99.7% accuracy on the training set and 98.7% on the held-out validation set. Users can retrain the network on their own labeled data; network weights are stored in standard MATLAB deep-learning format and the training script is provided with the software.

### Registration initialization using point clouds

Because B-spline deformable registration is a non-convex problem, robust initialization is critical. LightSuite first establishes a global similarity alignment by matching structural point clouds derived from the sample and from the Allen Common Coordinate Framework (CCF) v3 atlas template. The atlas template, natively at 10 µm, was downsampled to the 20 µm registration resolution, and the registration volume was intensity-normalized (divided by twice the 99.9th percentile of a central sub-volume) before cloud extraction.

#### Sample point cloud

The 20 µm registration volume was divided into contiguous 3D tiles, each spanning approximately one-third of the volume’s longest dimension along every axis. Within each tile, a background threshold was computed as the larger of (a) twice the 5th percentile of a random subsample of up to 10,000 local voxel intensities and (b) the global intensity mode of the entire volume. A structural saliency map was computed as the local 3D image gradient magnitude divided by voxel intensity. Voxels that exceeded both the background threshold and a saliency ratio of 5 were retained as candidate points. Points within a thin margin of the volume boundary were discarded to prevent edge artifacts. The aggregated point cloud was then randomly downsampled to 10% of its size and passed through a 3D point-cloud denoising step to yield a refined representation of the sample anatomy.

#### Atlas point cloud

The CCFv3 atlas template was downsampled to 20 µm, and voxels outside the brain annotation mask were set to zero. Saliency was computed as the 3D gradient magnitude divided by a Gaussian-smoothed version of the atlas volume (σ = 20 voxels). Voxels whose saliency exceeded 5 and whose intensity exceeded twice the 5th percentile of a sampled subset were retained. Unlike the sample cloud, the atlas cloud was used at full density without random downsampling or denoising.

#### Similarity alignment

Both point clouds were downsampled with a non-uniform grid filter before fitting. A similarity transformation (rotation, uniform scaling, translation) was estimated using Bayesian Coherent Point Drift^35^ (BCPD), with outlier ratio 0.01, γ=1, and convergence tolerance 10^-8^. After a first-pass fit, point pairs whose nearest-neighbor distance exceeded 500 μm in either direction were discarded as structural outliers, and BCPD was rerun on the mutually consistent inlier set to finalize the similarity transform. Fitting a similarity rather than a rigid transform matters because tissue-clearing methods can introduce isotropic shrinkage or swelling that varies across samples.

#### Automatic correspondence matching

To seed the subsequent affine and deformable steps with anatomical correspondences, candidate point pairs between the sample and atlas clouds were identified automatically. Both clouds were grid-downsampled, the sample cloud was mapped into atlas space using the similarity transform, and mutually distant points (nearest-neighbor distance > 50 voxels in either direction) were removed. A non-rigid BCPD fit (affine + non-rigid; outlier ratio 0.01, γ=0.1, λ=3, β=15, convergence tolerance 10⁻⁶, with common normalization) was then computed solely to establish correspondences: for each sample point, its 40 nearest warped-atlas neighbors were examined, and a pair was retained only if the closest neighbor lay within 20 voxels and the median distance over the 40 neighbors was below 50 voxels. The retained sample points were transformed back to native sample space and stored, together with their matched atlas points, as the automatically detected corresponding points used downstream.

### Registration refinement with user-defined control points

After automatic initialization, users optionally add corresponding landmarks through a graphical interface that shows a sample slice (coronal, sagittal or axial) beside the matching atlas slice, with atlas annotation boundaries overlaid on the sample to indicate alignment quality. The user clicks alternately on the two images to place paired landmarks. Active learning is built into this overlay: whenever the sample and atlas landmark counts on a view match, a 3D affine transform is refit in real time from all pairs accumulated across views, and the atlas boundaries are re-warped and redrawn. The overlay therefore sharpens as points are added, so each new landmark is placed against a more accurate reference and residual mismatches become easier to see; a running mean-squared error and landmark count track progress. To spread coverage, candidate views are sampled at evenly spaced positions along all three cardinal axes over the central ∼90% of the volume in randomized order, and each view exposes only one quadrant of a 2×2 subdivision of the slice to concentrate placement locally.

### Affine and deformable registration

User-defined control points are combined with the automatically detected correspondences. To avoid redundancy, automatic points lying within 1 mm of any user point were discarded, and the remaining automatic points were thinned against one another at the same 1 mm spacing (both distances were tripled to 3 mm when point augmentation was disabled). A 3D affine transform was then fitted to the combined point set by least squares, relating atlas and sample coordinates, and applied to warp the atlas template into the sample frame. The affine-transformed atlas template was registered to the sample volume by B-spline free-form deformation in Elastix v5.1.0^74^, using a multi-metric, multi-resolution scheme with a recursive cubic B-spline transform and a final control-point grid spacing of 0.64 mm. The objective combined Advanced Mattes mutual information (48 histogram bins; weight 1.0) with a corresponding-points Euclidean-distance term (weight 0.2) that drew the user-defined atlas points toward their matched sample points. Optimization used adaptive stochastic gradient descent over four resolution levels. When no user landmarks were available, the pipeline reverted to an automated mode that used the thinned automatic correspondences for the point term at half the control-point weight; the point term was dropped entirely only when no correspondences of any kind existed. The forward (atlas-to-sample) B-spline transform and its numerically computed inverse (sample-to-atlas) were both stored as Elastix parameter files.

### Spinal cord volume preprocessing and registration

Spinal cord samples may be curved and twisted in a sample-specific manner that is incompatible with direct atlas registration. LightSuite first maps the cord into a linearized reference frame and then registers it to a spinal-cord atlas^49^.

#### Sample preparation

The multichannel volume was resampled to the 20 µm registration resolution, and the long (rostrocaudal) axis was identified as the largest volume dimension and moved to the third axis. The registration channel was binarized and morphologically dilated, and the largest connected component was retained as the cord mask. The rostrocaudal orientation was inferred from the cross-sectional area profile and the volume flipped to a consistent rostral-to-caudal direction when necessary. Slices whose cross-sectional area greatly exceeded the median (indicating attached brain tissue) were detected and excluded from the analyzed range.

#### Centerline annotation

A custom GUI displays each axial slice of the registration volume. On slices of their choosing, the user marks up to three anatomical points: the dorsal and ventral extremes of the cord and the central canal (left, right, and middle mouse clicks, respectively). Per slice, the canal position defines the cord center (or, when the canal is unmarked, the midpoint of the dorsal and ventral points), while the dorsoventral vector defines the cord orientation angle and radius. Side panels display the fitted center position (x, y) and orientation angle (θ) against slice index for continuous quality control, and the interface highlights the largest unannotated gap to guide sampling.

#### Smoothing-spline fit

The sparse, possibly noisy per-slice observations of center-x, center-y, orientation angle, and radius are each converted into a continuous trajectory along the rostrocaudal axis by solving the penalized least-squares system (*W* + *λL*)*c* = *b*, where *W* is a diagonal indicator matrix selecting annotated slices, *b* holds the observed values, and *L* = *D*₂ᵀ*D*₂ is built from a second-difference operator that penalizes curvature in the trajectory (with first-difference end conditions). Angular observations were phase-unwrapped before fitting and the resulting trajectory re-wrapped to (−π, π]. The regularization parameter λ (default 5000 for both position and angle, adjustable in the interface) controls trajectory smoothness, with smaller values appropriate for more tortuous samples.

#### Per-slice straightening and registration

For each axial slice, a rigid 2D transform was constructed from the fitted trajectory by translating the fitted canal position to the image center, rotating by the fitted orientation angle to standardize the cord pose, and translating to the center of the spinal-cord atlas. Applying these per-slice transforms produced a straightened volume. The atlas was initialized into this volume by a z-axis scaling and translation matching the annotated extent, refined by an Elastix affine registration driven by Advanced Mattes mutual information, and finally registered with the same multi-metric B-spline free-form deformation used for brain volumes (final control-point grid spacing 960 μm), combining mutual information with a landmark term derived from user-defined correspondences between sample and atlas structures.

### Quantification of cell counts and intensities across samples

All quantification was performed in Allen CCFv3 atlas space after registration, at the substructure level of the Allen parcellation unless stated otherwise. Detected somata were carried into atlas space through the stored sample-to-atlas transform and assigned to the annotation label of the voxel in which they fell; detections lying outside the brain annotation or beyond the volume bounds were discarded. Cells were split by the midplane into the two hemispheres. For every substructure we counted the detections it contained and measured its volume as the number of annotation voxels multiplied by the voxel volume (10^-6^ mm^3^ per voxel). Regional cell density was defined as the count divided by the regional volume. Because unilateral illumination and dissection can affect the two sides differently, counts and volumes were computed separately per hemisphere and then averaged across hemispheres, so that each region contributes a single per-hemisphere count, volume and density per animal. In parallel, each animal’s registered signal volume was resampled into atlas space, and for every substructure and hemisphere we took the median voxel intensity within the region as its fluorescence signal and the median intensity of the out-of-brain voxels as that hemisphere’s background. Intensities are reported relative to this background. When values were aggregated from finer to coarser parcellation levels (substructure → structure → division), counts were summed, intensities were combined as volume-weighted means, and the two hemispheres were averaged throughout.

#### Region masking and sample quality control

We restricted all analyses to gray-matter regions, excluding fiber tracts, the ventricular system, white matter, nerves and unassigned labels. Regions with unreliable signal were treated as missing and omitted from an animal’s statistics: within each animal a region was masked when its background-relative intensity fell below a fixed threshold (0.5) or more than two robust standard deviations below the animal’s median across regions, and regions with a density below 10^-2^ cells/mm^3^ were likewise treated as missing. At the whole-sample level, we used consistency with the cohort as an expression-based quality criterion: a brain whose regional density profile did not correlate with the common expression axis shared across animals (see below; Spearman ρ ≤ 0.5) was considered to lack activity-dependent labeling and was excluded from cross-animal averaging and map building (1 of 34 TRAP2 brains).

#### Cross-animal normalization

Raw cell counts and intensities vary across animals with labeling efficiency and imaging conditions. To place samples on a common scale, we estimated a single non-negative axis of common expression by rank-one non-negative matrix factorization of the regional-value matrix across samples, restricted to substructures with complete data in every animal within a predefined set of major divisions (isocortex, cortical subplate, hippocampal formation, striatum, pallidum, thalamus, hypothalamus and midbrain). The factorization returns one non-negative weight per animal, which we divided by its median across animals to obtain a per-animal normalization factor. Regional densities or intensities were divided by this factor before cross-animal averaging, so that samples differing only in overall labeling strength were brought into register; densities and intensities were normalized independently, each from its own factorization. The same rank-one factorization, applied to per-region soma diameters, provided the multiplicative normalization used to compare diameters across animals. Laminar profiles of soma diameter (Fig. 4L) were estimated with a linear mixed-effects model fitted to the normalized per-area, per-layer diameters of all animals, with cortical layer as a categorical fixed effect and a random intercept for each cortical area (diameter ∼ layer + (1|area)). The model was fitted separately for LightSuite and brainreg registrations, and the reported profiles and 95% confidence intervals are the fixed-effect predictions for each layer, with area-specific offsets removed.

#### Hemispheric asymmetry

To quantify hemispheric asymmetry in intensity or cell counts, we defined an asymmetry index for each atlas substructure as (*S_L_* − *S_R_*)⁄(*S_L_* + *S_R_*), where *S_L_* and *S_R_* are the values in the left and right hemispheres. For each brain, we report the asymmetry index as a median across substructures (Fig. 3J).

#### Brain-wide maps and cortical flatmaps

To visualize expression across the cohort, we built a brain-wide cell-density map by placing every retained detection at its atlas voxel, accumulating across all quality-controlled light-sheet brains with a per-animal weight equal to the inverse normalization factor, averaging the two hemispheres, and smoothing with a 3D Gaussian (σ = 35 µm); the accumulated counts were divided by the voxel volume to yield densities in cells/mm^3^. An analogous intensity map was formed by averaging the background-relative signal volumes in atlas space across animals, weighted by the inverse of the intensity normalization factor and hemisphere-averaged. For cortical visualization, the averaged density map was projected onto the Allen cortical-surface flatmap separately for each cortical layer with the ccf_streamlines package (Allen Institute, https://github.com/AllenInstitute/ccf_streamlines), which implements the Allen CCFv3 cortical flatmap^16^, with atlas area boundaries overlaid, and read out along cortical depth.

### Slice module: volumetric reconstruction from coronal sections

LightSuite includes a module that reconstructs serial sections acquired by conventional wide-field microscopy into a volume in atlas space and registers them to the Allen CCFv3.

#### Preprocessing and assembly

Each section was median-filtered (3×3), and downsampled to a common processing resolution (5 μm for intensity analysis and 1.25 μm for cell detection). Empty (zero) pixels from tiling were replaced by the 1st-percentile intensity of the section, and for DAPI channels an optional local standard-deviation filter was applied to enhance nuclear contrast. Sections were cropped to the tissue extent and symmetrically padded to a uniform field of view. Using an annotation GUI, users can correct section ordering, flip individual sections to a consistent orientation, and flag heavily damaged sections (e.g., folding) for exclusion.

#### Initial volumetric alignment

The registration channel of the assembled stack was converted into a per-section 2D point cloud of high-signal pixels, and the sections were spaced along the rostrocaudal axis according to the physical section thickness to form a 3D sample cloud. A corresponding cloud was extracted from the atlas template. The atlas and sample clouds were then aligned by an iterative optimization that alternated between two rigid fits: a global 3D rigid transform from atlas to sample space, estimated by Coherent Point Drift on the whole cloud, and a set of per-section in-plane 2D rigid transforms, estimated by iterative closest point (ICP) between each section and the atlas plane at its rostrocaudal position. In each pass, the current per-section transforms were applied to the sample cloud before re-estimating the global 3D transform, and the atlas cloud was then re-warped by that transform before re-estimating each per-section transform; three passes were run. This procedure yielded both an initial placement of the whole stack within the atlas (the global 3D rigid transform) and a per-section in-plane correction that straightens each section within the assembled sample volume.

#### Cutting-angle determination

Because serial coronal sections are rarely cut exactly orthogonal to the rostrocaudal axis, a dedicated GUI lets the user rotate the 3D atlas interactively until the displayed atlas plane matches a given section, saving the corresponding camera viewing direction for that section. The saved per-section viewing directions were averaged into a single mean cutting normal, which was used to correct the rotation component of the global atlas-to-sample transform (about the atlas center) so that atlas planes are resampled at the true mean cutting angle.

#### Per-section registration

For each section, the matching atlas plane was extracted at the section’s assigned atlas index. Users could place corresponding landmarks between the section and the atlas plane through a GUI; when at least five landmark pairs were available, a 2D affine transform was fit directly from them, otherwise an affine was estimated automatically from image-derived point clouds. The affine-aligned atlas plane was then registered to the section by 2D B-spline free-form deformation in Elastix, using the same multi-metric scheme as for volumes (Advanced Mattes mutual information plus a corresponding-points term, 32 histogram bins, 0.96 mm final grid spacing). Both the forward (atlas-to-section) and inverse (section-to-atlas) transforms were stored per section.

#### Volumetric reconstruction

To build a volume in atlas space, each section’s channels were mapped through the stored inverse B-spline and inverse affine transforms into the atlas coordinate frame and placed at the section’s assigned rostrocaudal index. Gaps between sections were filled by nearest-neighbor interpolation along the rostrocaudal axis, and the resulting volume was warped into the full atlas space using the global 3D rigid transform. Per-section cell detection used the same 2D adaptation of the SBR-based algorithm described above, and detected coordinates were carried into atlas space through the same stored transforms.

### Registration benchmarking

LightSuite registration was benchmarked against brainreg^18^ version 1.0.10, an automated tool based on mutual-information image registration. We used the same 20-µm volumes used by LightSuite and registered them to the 25 μm Allen Atlas with default parameters.

Because expert-drawn region-of-interest overlap metrics are sensitive to segmentation choices, we instead used a set of automated, reproducibility-based metrics computed on gray-matter substructures of the Allen atlas, and excluding fiber tracts, ventricular structures and unassigned regions. For each animal, regional tdTomato+ cell densities were obtained as counts divided by regional volume, and regional fluorescence intensities as volume-weighted means, after masking regions with unreliable background signal. We then computed, using Spearman correlations throughout: (i) pairwise correlations of regional intensity profiles between every pair of independently registered animals; (ii) the same pairwise correlations using regional tdTomato+ density profiles; and (iii) the correlation between each animal’s regional tdTomato+ density and a standardized anatomical density profile (total neuronal density). The pairwise correlations were evaluated both across the full set of substructures and separately within each coarse anatomical division, to test consistency at finer spatial granularity. As a further reproducibility measure, each animal’s density profile was correlated with a common profile extracted across all samples by non-negative matrix factorization. For every metric, higher values indicate that analogous anatomical features were placed more consistently into the same atlas coordinates; all metrics were computed identically for LightSuite and brainreg to allow direct comparison.

To evaluate registration accuracy against anatomical ground truth, layer 4 cortical flatmaps centered on the primary somatosensory barrel cortex were extracted for registered samples and the CCFv3 template. A 2D rigid transformation (rotation about the crop center and translation) was manually fitted to align each sample’s barrel field to the reference template. Registration error was quantified as the root-mean-square per-pixel displacement across the barrel field in micrometers (incorporating both translation and rotation), along with the translation of the crop center and rotation magnitude. A hemisphere was included when its barrel pattern could be resolved in the layer 4 flatmap after LightSuite registration (n=18 hemispheres); hemispheres with tissue damage or insufficient autofluorescence contrast over the barrel field were excluded. For the comparison with brainreg, hemispheres whose barrel field could not be resolved after brainreg registration were additionally excluded (7 of 18).

### Cell detection benchmarking

We compared LightSuite against Cellpose v4.0.8^24,75^ on field-of-view crops (1000×1000×32 voxels) extracted from four cleared brains. Ground-truth cell centers were annotated manually using a dedicated interface that displays maximum-intensity projections of consecutive depth blocks; each click was snapped to the local intensity maximum within a small 3D neighborhood, yielding a set of validated 3D cell positions per volume. LightSuite detection was run with and without the artifact-rejection classifier. Cellpose was run in 3D mode on the GPU (anisotropy factor 3.47, intensity normalized to the 90th–99.99th percentile) and applied to the raw volumes; Cellpose cell centers were taken as the median voxel coordinate of each segmentation mask.

Detected and annotated cell centers were matched one-to-one in physical space by minimum-total-distance bipartite assignment, forbidding pairs farther apart than 10 µm; from the resulting matched pairs we computed sensitivity, precision, and their harmonic mean (F1 score) for each method. Sensitivity (recall) was defined as the number of matched pairs divided by the total number of annotations, and precision as the number of matched pairs divided by the total number of detections (set to zero when a method produced no detections). We additionally recorded the number of detected cells and the processing time per volume, which differed by roughly two to three orders of magnitude between the two approaches. As a label-free quality measure, we extracted a fixed voxel neighborhood (18×18×12 voxels) around every detected center and computed the pairwise correlations between these neighborhoods across all detections within a volume; a higher median correlation indicates that detections more consistently coincide with stereotyped cell-like intensity profiles. All metrics were computed identically across methods.

### Cylindrical implant localization

Implants were annotated on the 20 µm registration volume by marking points on, within and around the implant on every coronal slice where it was visible; these were mapped into Allen CCFv3 space^16^ using the registration transform, and fitted at 10 µm resolution. The implant trajectory is defined as the line minimizing the radius of the cylinder enclosing 95% of the points, evaluated as the mean of the ordered point-to-line distances between the 80th and 95th percentiles. Implants enter approximately perpendicular to the brain surface, so the axis was constrained to within 45° of the local surface normal vector about each transverse axis; that normal was obtained by fitting a plane to the labelled boundary voxels of the annotation volume within 600 µm of the entry point, requiring open space beyond the patch so that fissure and ventricle walls are excluded. The constrained problem was solved by sequential quadratic programming over the two tilt angles and the two lateral coordinates of the line, with a rotation-invariant regularization toward the surface normal that is appreciable only for annotations too shallow to determine the tilt. The implant bottom face was placed at the 99th percentile of the axial coordinate of the enclosed points, and parcellation labels and fluorescence beneath the implant were sampled on square segments twice as large as the nominal implant radius perpendicular to the axis, at the face and at 100 µm intervals to 400 µm below it.

### Functional ultrasound imaging

fUS volumes measure cerebral blood volume at coarse spatial resolution, and their contrast reflects vasculature rather than the cytoarchitecture annotated by the Allen CCF, so we first refined an existing vascular registration template^21^. The high-resolution power-Doppler template distributed with the dataset (50 µm isotropic) was registered into the Allen CCF v3 with the same point-cloud-initialized, multi-metric B-spline pipeline used for cleared brains, and the Allen annotation was transferred onto the warped vascular template. The template was symmetrized across the midline to remove residual left–right asymmetry. The resulting vascular template and its inherited annotation served as the registration target for all fUS sessions.

To illustrate registration performance, we used previously published fUS data from anesthetized mice^59^. We refer the reader to the original publication for acquisition and analysis details regarding the power-Doppler fUS data, and the block-design visual-object paradigm (alternating 12 s of gray and 12 s of images, with object and texture-scrambled blocks interleaved). Activation and object-preference maps were computed as described below.

Because each imaging session is positioned freehand, registration proceeds in two stages. First, repeated recordings of a given mouse are aligned to a common within-mouse template. To build the within-mouse template, a motion-robust volume is first distilled from each recording. Ten candidate volumes are formed, each the median of 50 frames drawn at random from the middle 80% of the recording; every frame is compared against these candidates, and the template is refined from the most mutually consistent frames, those in the top 10% of agreement, so that movement-corrupted frames are suppressed. Every session is resampled to the 50 µm atlas resolution and rescaled to a common intensity so that no single recording dominates the average. One recording is then designated as the reference, or seed, that defines the animal’s common anatomical space, and each remaining session is rigidly aligned to it by mutual-information registration. Second, the template is registered to the atlas. Its orientation relative to the atlas axes is established first, and a set of matching anatomical landmarks, at least five and spanning all three dimensions, is placed by hand in the template and in the atlas to anchor the alignment. Affine and cubic B-spline registrations follow, each driven jointly by image mutual information and by the agreement of the manually matched landmarks. The cubic B-spline fit is constrained by a grid of 1.6 mm isotropic spacing. The resulting transformation, together with its inverse, defines a bidirectional mapping between each mouse’s anatomical space and the atlas, and overlays of the warped parcellation on the template are retained for visual quality control.

Activation maps were computed for each session in the mouse’s native space and then carried into atlas space with the transforms above. Visual responses were estimated with a general linear model in which a single boxcar regressor covered every stimulus block; the t-statistic of that regressor is the visual t-score (Fig. 7C). Voxels whose group visual t-score exceeded 1 were defined as visually responsive, and all object measures below are reported within that mask.

Object preference was estimated from a second general linear model fitted to the same data, with two regressors: one for the object blocks and one for the texture-scrambled blocks. To isolate the transient part of the response, each regressor covered only the first 6 s of each 12-s block. The object contrast of a session is the difference between the two fitted weights, β_object_ − β_scrambled_. For each mouse, we divided the mean of this contrast across sessions by its standard error across sessions, giving a per-mouse object t-score at every voxel; voxels covered by fewer than two sessions were left undefined. The group object t-score map (Fig. 7E,G) is the one-sample t-statistic of these per-mouse values across the seven mice. Where the object response is summarized as an index rather than as a contrast (Ext. Data Fig. 8G,H), we used the visual object sensitivity index, *VOSI* = (*O* − *S*)/(|*O*| + |*S*|), where *O* and *S* are the areas under the baselined, trial-averaged response to object and scrambled blocks over the 12-s stimulus window; *VOSI* was computed per session and averaged across sessions. To compare object preference across the area groups (Fig. 7H), we used the atlas area rather than the voxel as the unit of replication, because neighboring voxels are spatially correlated and the groups differ about tenfold in voxel count. For each mouse, the transient object t-score was summarized as the median over the visually responsive voxels of each atlas area with at least 10 such voxels (16 areas). These area-by-mouse values were fitted with a linear mixed-effects model with area group as a fixed effect and crossed random intercepts for mouse and area (y ∼ group - 1 + (1|mouse) + (1|area)). With one coefficient per group, the model gives the population mean and 95% confidence interval of each group; each group mean was compared with zero by a Wald test, with Benjamini-Hochberg correction across the four groups. The omnibus group effect was assessed with a likelihood-ratio test between maximum-likelihood fits of y ∼ group + (1|mouse) + (1|area) and y ∼ 1 + (1|mouse) + (1|area).

### Software, computing hardware and runtime

All processing was performed on a workstation equipped with an Intel Core i9-10900X CPU (3.70 GHz), 64 GB DDR4 RAM, and an NVIDIA RTX A4000 GPU (16 GB VRAM). Whole-brain cell detection required 3–4 h, automated point-cloud registration 5–6 min, and interactive landmark curation 10–20 min (∼100 landmarks). LightSuite is implemented in MATLAB, and all analyses reported here were run in MATLAB R2024a (MathWorks). The exceptions are the cortical flatmap projections, which use the Python package ccf_streamlines, the Python-based tools used for benchmarking (brainreg, Cellpose), and the Elastix and BCPD binaries, which LightSuite calls externally.

### Statistics

All statistical tests were two-sided unless specified otherwise. Wilcoxon signed-rank tests were used for paired non-parametric comparisons; Spearman rank or Pearson correlation coefficients were computed as indicated. Sample sizes (n) denote independent animals, sample pairs, or anatomical parcels. Error bars represent robust standard deviations, s.e.m., or 95% robust confidence intervals as detailed in figure legends.

## Data and code availability

LightSuite source code is openly available at github.com/dimokaramanlis/LightSuite under a GPL-3.0 license. Example datasets will be made available in Zenodo (https://doi.org/10.5281/zenodo.21338217) and a Python version of the registration module will become available at github.com/Wyss-Microscopy/PyLightSuite.

## Author contributions

DK and SEB conceived the study. DK developed and implemented LightSuite, performed the analyses and prepared the initial manuscript draft. DK, YX, CGF, MA, AM, ID and SG collected, processed and imaged samples. LAFP implemented the Python version of LightSuite. JMMdP and EM provided the functional ultrasound data and assisted in interpreting the results. SEB supervised the research, secured funding, provided resources and contributed to the design and interpretation of the study. DK and SEB wrote and revised the manuscript with input from all authors.

## Acknowledgements

We thank Laura Batti and members of the Advanced Microscopy Facility at the Wyss Center in Geneva for technical assistance during light-sheet imaging. SEB and SG were supported by the Swiss National Science Foundation (Project numbers: 320030-231711 and PZ00P3-208875). JMMdP and EM were supported by the Else Kröner Fresenius Foundation through the Else Kröner Fresenius Center for Optogenetic Therapies, the Niedersächsisches Ministerium für Wissenschaft und Kultur through zukunft.niedersachsen, and the Deutsche Forschungsgemeinschaft (DFG, German Research Foundation) through Germany’s Excellence Strategy EXC 2067/1-390729940, the Project number 537609931 (FOR 5807, project 1) and the Collaborative Research Center 1690 (Project B06).

## SUPPLEMENTARY FIGURES

**Ext. Data Fig. 1.**
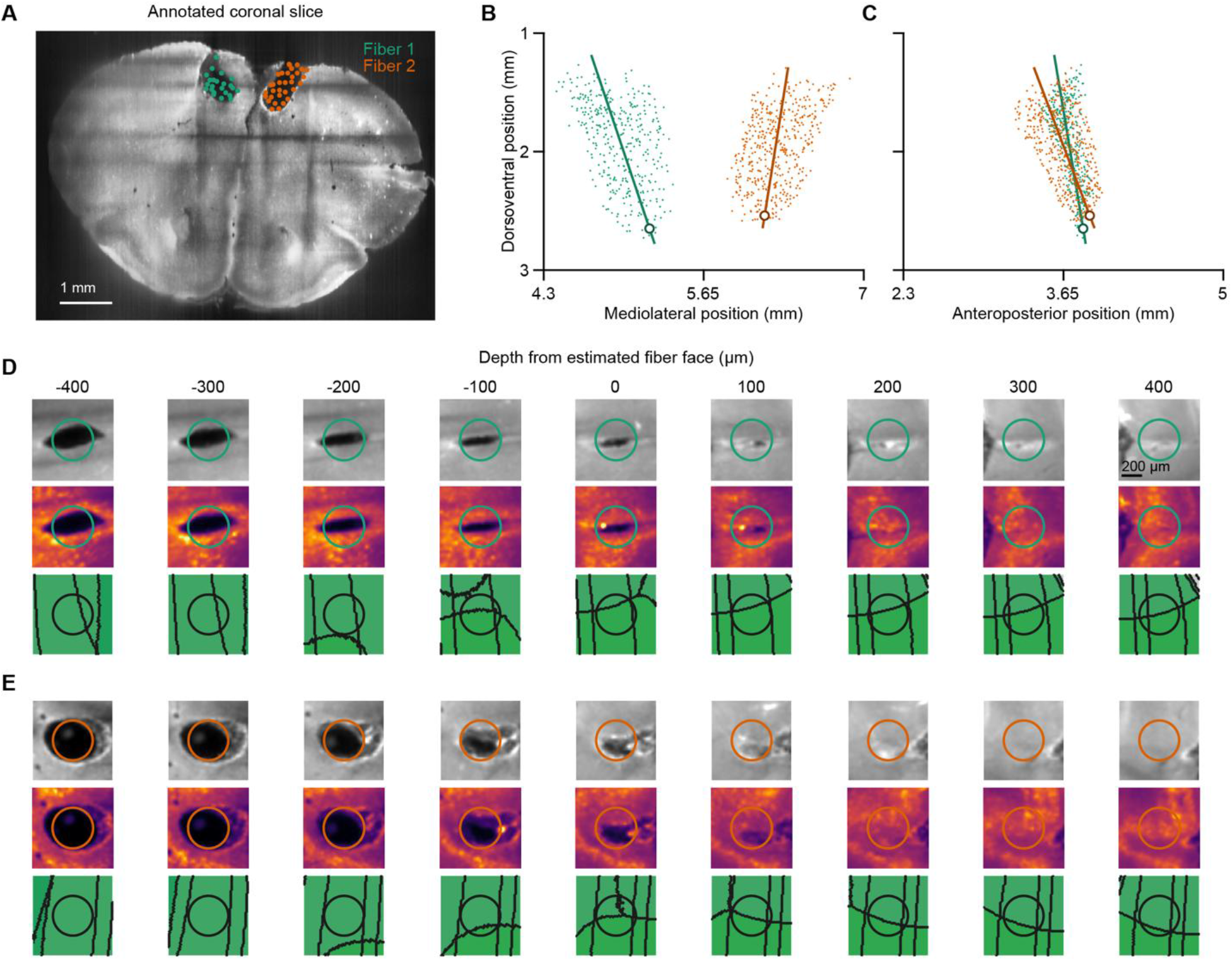
3D reconstruction of optic fiber implant trajectories. **A**) Example coronal slice from an implanted mouse brain imaged at 488-nm for autofluorescence with light-sheet microscopy. On top are points added by the user to cover the damage left by two 400-μm-diameter optic fiber implants. **B**) All annotated points (fiber 1: 369 points; fiber 2: 448 points) projected along the anteroposterior axis of the Allen Mouse Brain Atlas. Lines indicate the fitted center of each fiber, and white dots at the ends of the lines mark the estimated fiber face. **C**) Same as (B), but projected along the mediolateral axis. **D**) Top: projection of the autofluorescence volume around the fiber 1 track across nine different depths around the estimated fiber face. Note that the fiber damage on top of the face is aligned with the estimated diameter. Middle: same as top, for 561-nm excitation, which reports FusionRed expression from the viral injection. Bottom: the same projection of the Allen Mouse Brain Atlas annotation volume; lines indicate area boundaries and colors follow the Allen convention. **E**) Same as (D), but for fiber 2.

**Ext. Data Fig. 2.**
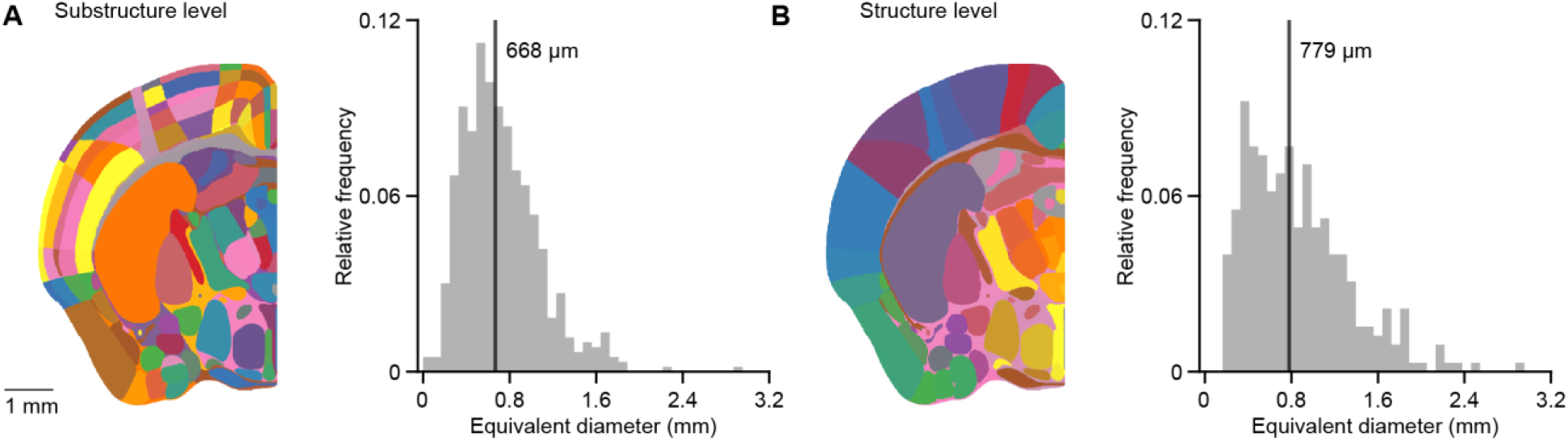
Spatial scale of mouse brain areas. **A**) Left: an example coronal slice of the Allen Mouse Brain Atlas parcellation at the substructure level. Right: distribution of the equivalent diameter (the diameter of a sphere with the same volume as each area) across substructures; the black line marks the median. **B**) Same as (A), at the coarser structure level.

**Ext. Data Fig. 3.**
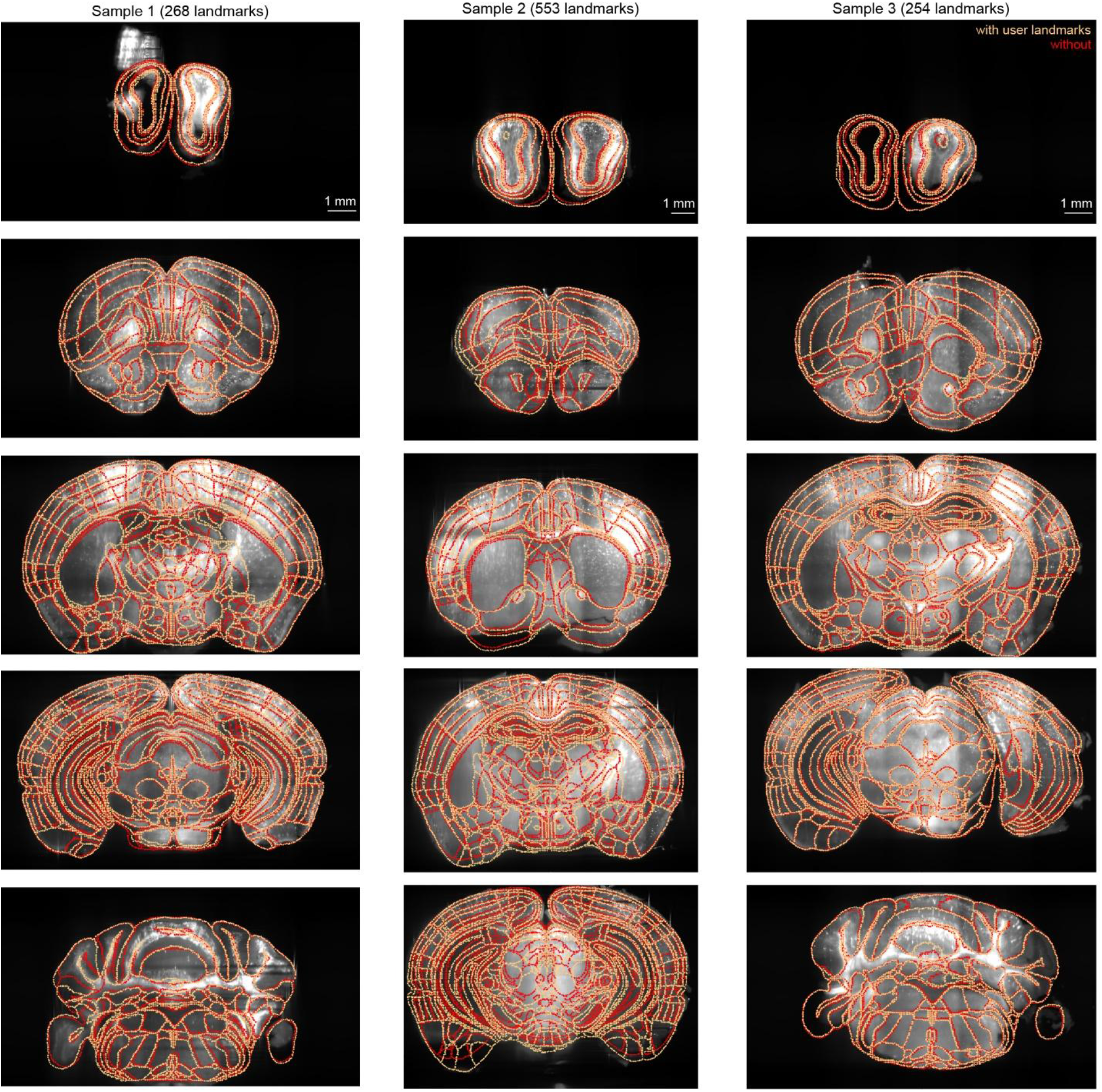
Comparison of non-rigid atlas registration with and without user-defined landmarks. Five representative coronal levels from each of three brains, showing B-spline deformable fits with user landmarks (gold) versus automated image-only registration (red). The number of user landmarks is given above each sample.

**Ext. Data Fig. 4.**
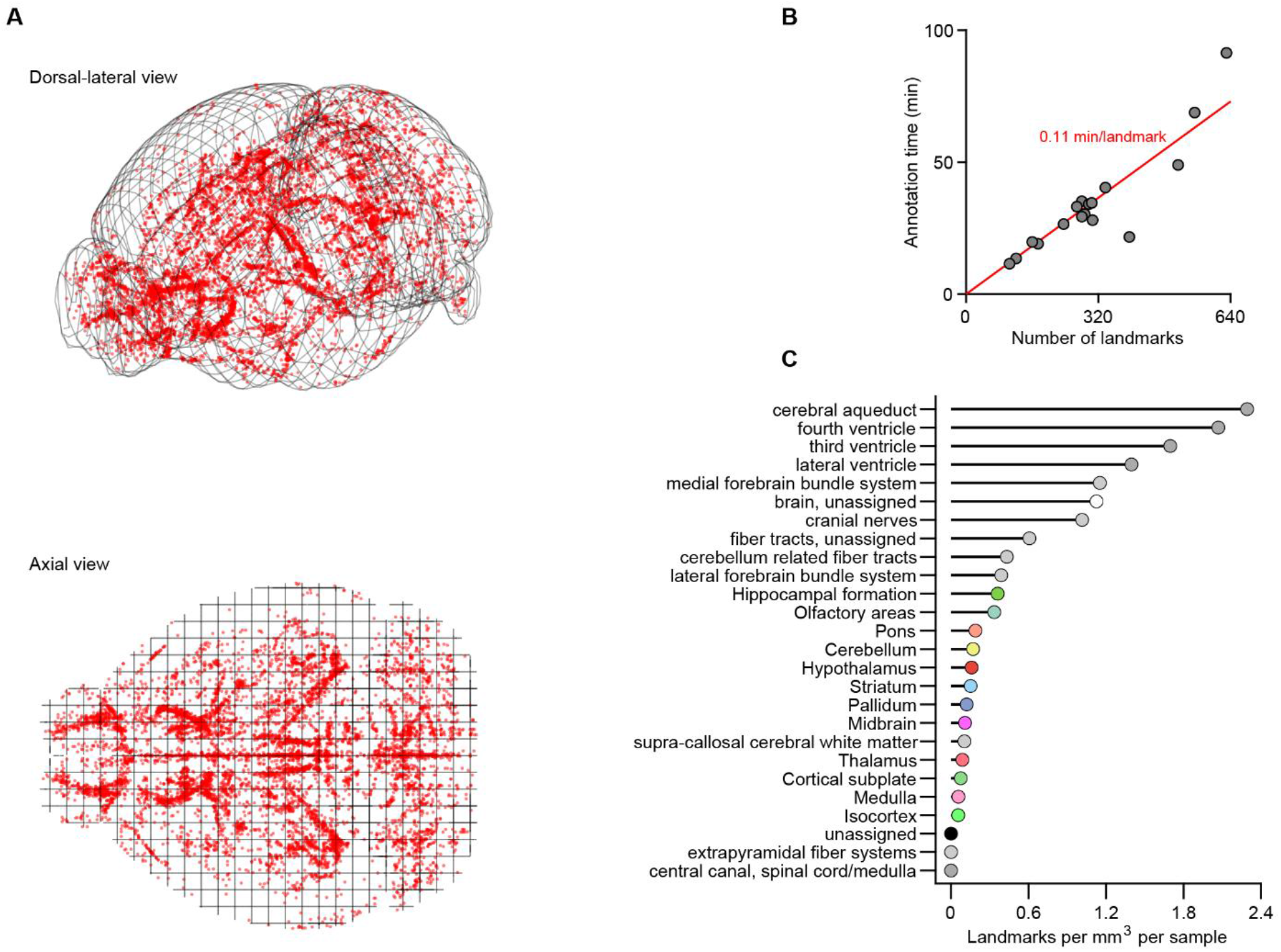
Spatial distribution and annotation timing of user landmarks. **A**) Distribution of user-placed landmarks (red dots) in CCFv3 space across 17 brains in dorsolateral (top) and axial (bottom) projections. Most points are placed internally and are symmetric across sides. **B**) Annotation time scales linearly with landmark count (Pearson’s r=0.91, p=3.7·10^-7^). Annotation time was computed as the accumulated inter-annotation interval, excluding gaps above 3 minutes (which likely reflect re-evaluation). **C**) Landmark density across Allen divisions; placement concentrates along ventricles, fiber tracts, hippocampus and olfactory bulbs.

**Ext. Data Fig. 5.**
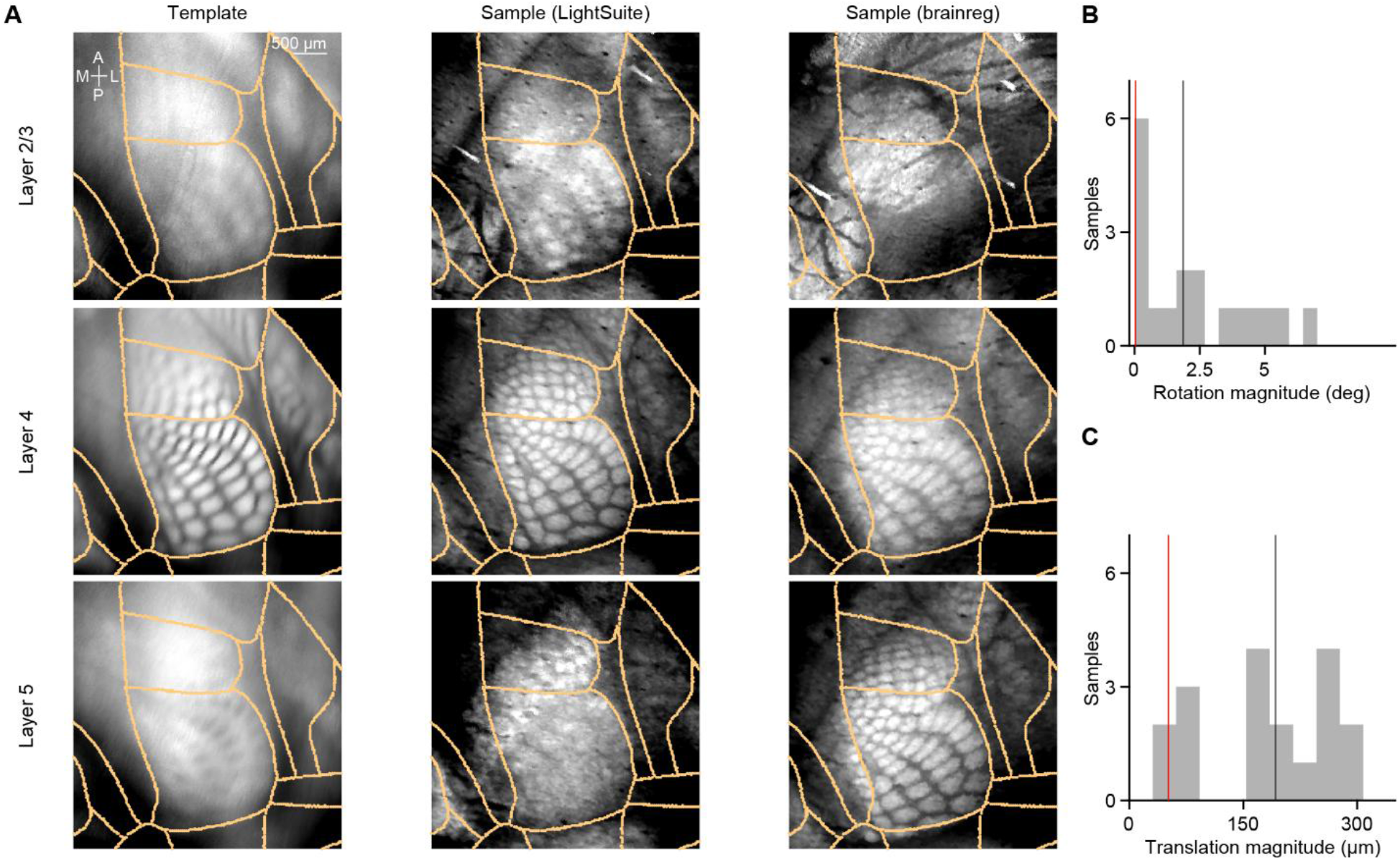
Barrel fields in registered brains. **A**) Left: Average flatmap projections of layers 2/3, 4, and 5 of the Allen Mouse Brain Atlas reference template. Middle: Layer-specific maximum projections of an autofluorescence sample registered with LightSuite. Right: Same sample registered with brainreg. **B**) Distribution of the rotation magnitude required to align sample barrel fields to the template (see Methods; red line indicates the sample in A; black line indicates the population median, n=18 hemibrains from 9 samples). **C**) Distribution of the translation magnitude for the same samples.

**Ext. Data Fig. 6.**
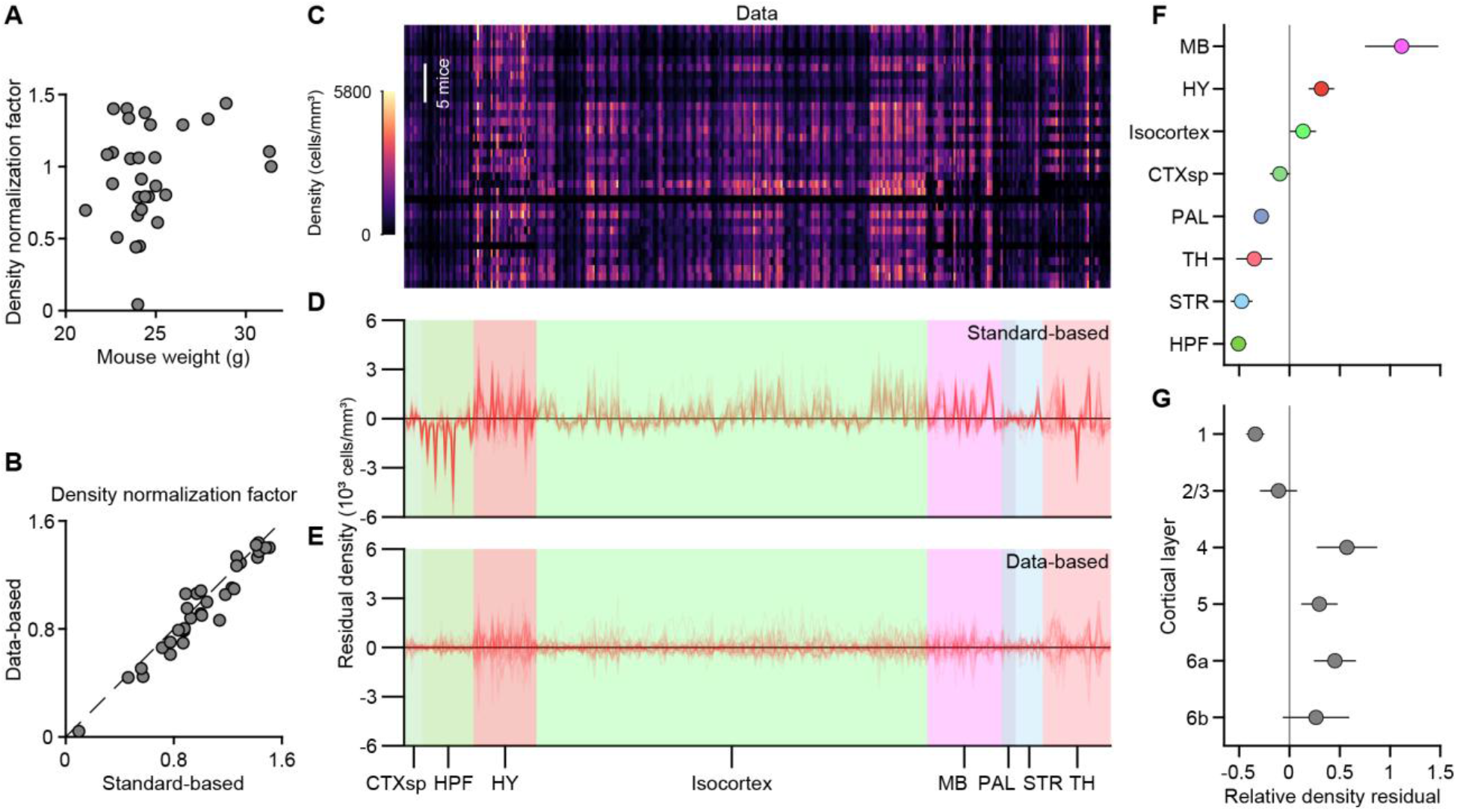
Cross-animal normalization and residual tdTomato+ activity profiles. A) Per-sample density normalization factor versus mouse body weight prior to 4-OHT injection (n=30 mice with recorded weight). The normalization factor is not significantly correlated with body weight (Spearman’s ρ=0.16, p=0.40), arguing against weight-dependent dosing dilution. **B**) Data-based normalization factor (derived from rank-one non-negative matrix factorization and used across all main figures) plotted against the standard-based factor (obtained by projecting measured regional cell densities onto the normalized anatomical density vector, i.e. the regional neuron densities of the standard cell atlas used in Fig. 4A). Points fall closely along the identity line (dashed line; R^2^=0.90). **C**) tdTomato+ cell density across brain substructures (columns) for each light-sheet volume (rows), ordered by Allen brain division. **D**) Residual cell density across substructures after subtracting a rank-one prediction based on the standard reference anatomical density profile. Individual sample traces are shown in red against Allen division color bands. Because tdTomato+ density departs from cytoarchitecture in a similar way across animals, these residuals retain a shared profile. **E**) Same as (D), but showing residuals after subtracting the data-driven rank-one non-negative matrix factorization model, which absorbs this shared profile. The black horizontal line marks zero residual. **F**) Relative density residual, (data−prediction)/prediction taken as the median across samples, aggregated per Allen division. Markers are area-volume–weighted means colored by Allen division; error bars are 95% CI. **G**) Relative density residual across cortical layers (area-volume–weighted mean ± 95% CI).

**Ext. Data Fig. 7.**
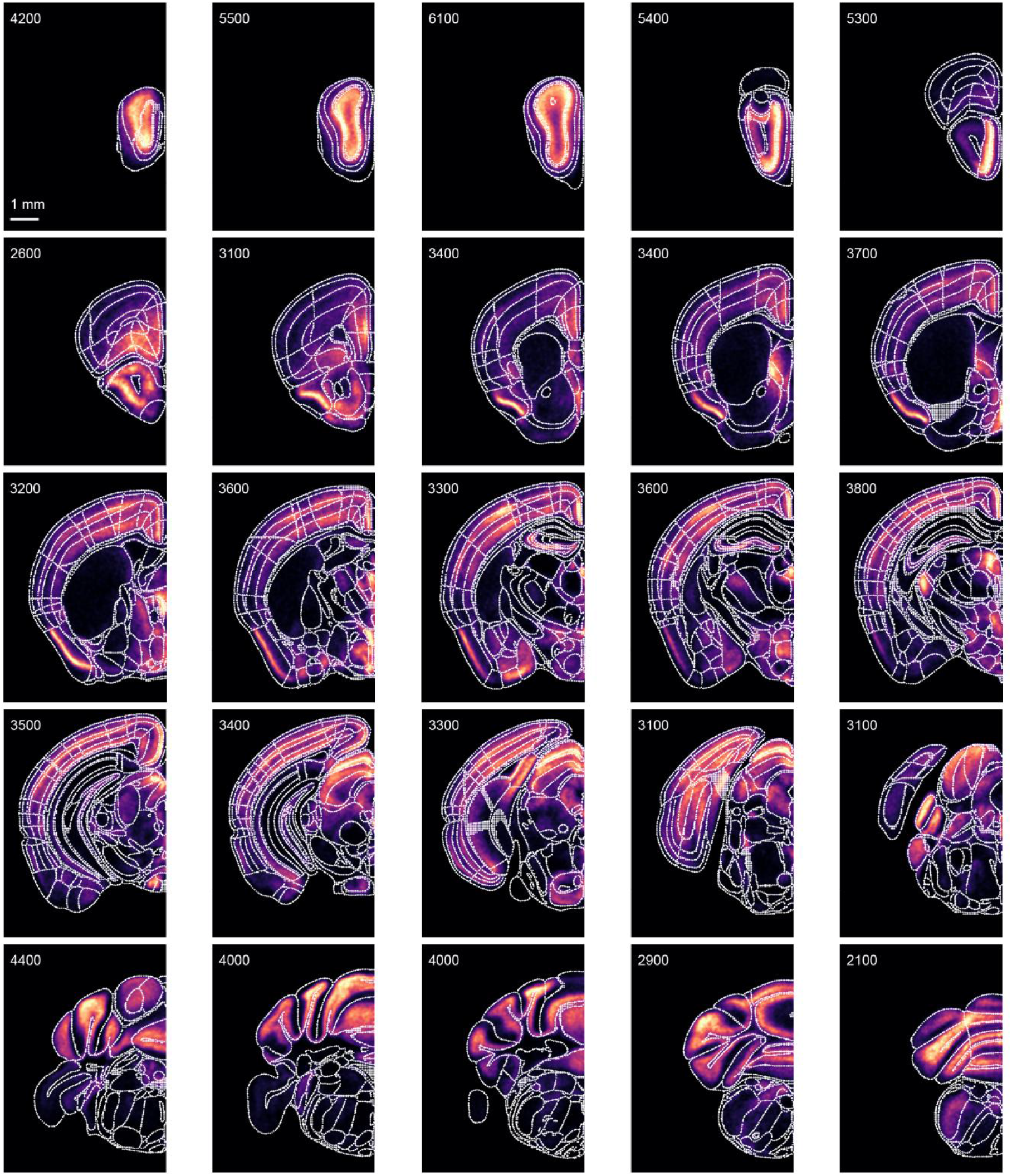
Brain-wide Fos activity atlas. Cohort-averaged tdTomato+ cell-density map (n = 33 brains) collapsed onto a single hemisphere across serial anteroposterior planes. For each plane illustrated, the number shown corresponds to the brightest value in the colormap in cells/mm^3^.

**Ext. Data Fig. 8.**
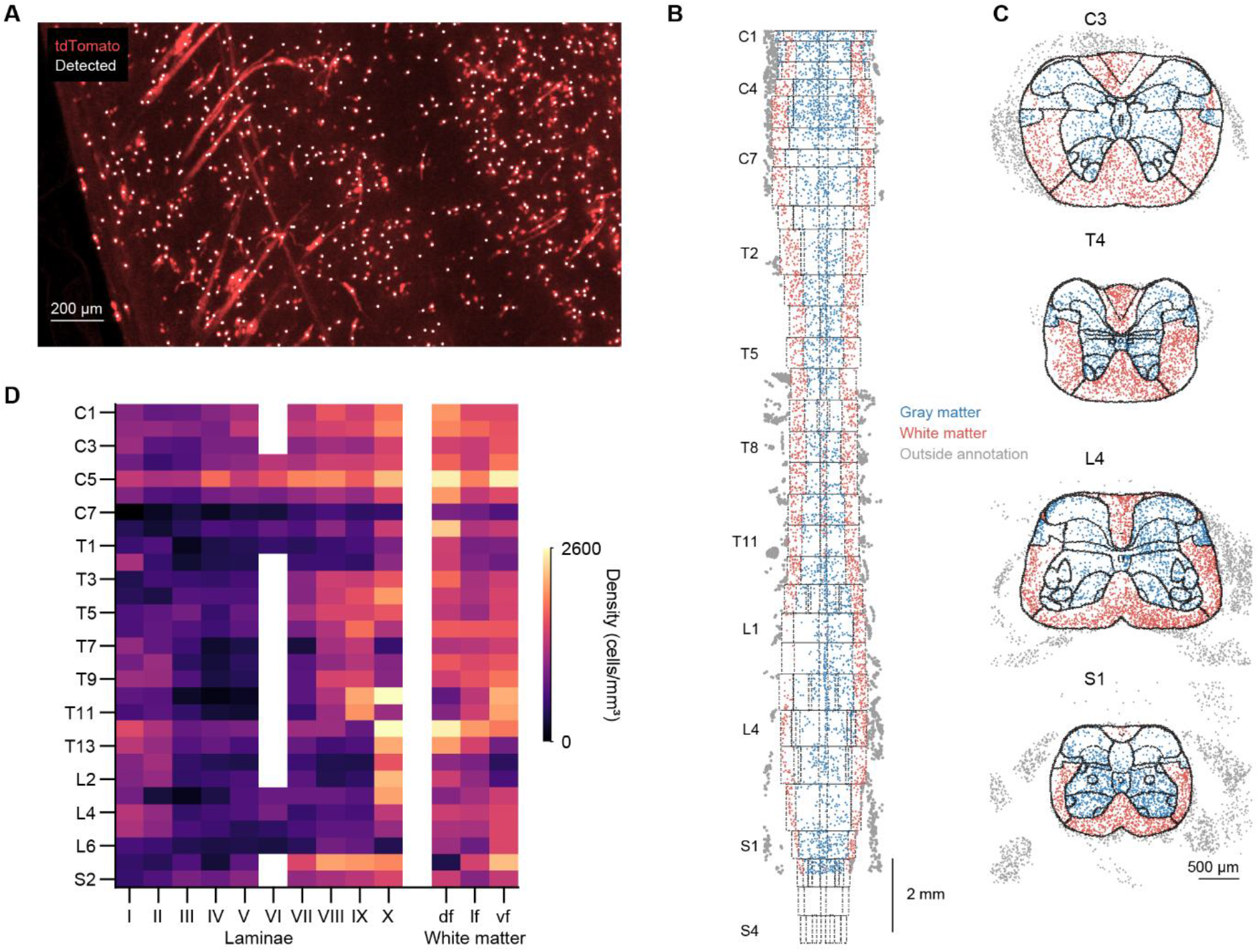
Quantification of oligodendrocyte-precursor-cell-derived (OPC-derived) tdTomato+ cell density in a spinal cord sample. **A**) Maximum-intensity projection of a representative raw light-sheet field (tdTomato, red) with LightSuite-detected OPC-derived tdTomato+ somata overlaid (white dots). **B**) Horizontal section through center of the registered spinal cord volume, shown from rostral (top) to caudal (bottom). Black dots mark atlas boundaries, and horizontal lines indicate segment borders. Overlaid points represent detected cells located within 50 µm of the section plane, colored by anatomical compartment: blue for gray matter, red for white matter, and gray for detections outside the atlas annotation. **C**) Four representative transverse sections at cervical (C3), thoracic (T4), lumbar (L4), and sacral (S1) levels with atlas boundaries (black) and detected cells colored as in (B). **D**) Heatmap of detected OPC-derived tdTomato+ cell density across spinal segments (rows) and anatomical structures (columns), divided into gray matter laminae and white matter funiculi.

**Ext. Data Fig. 9.**
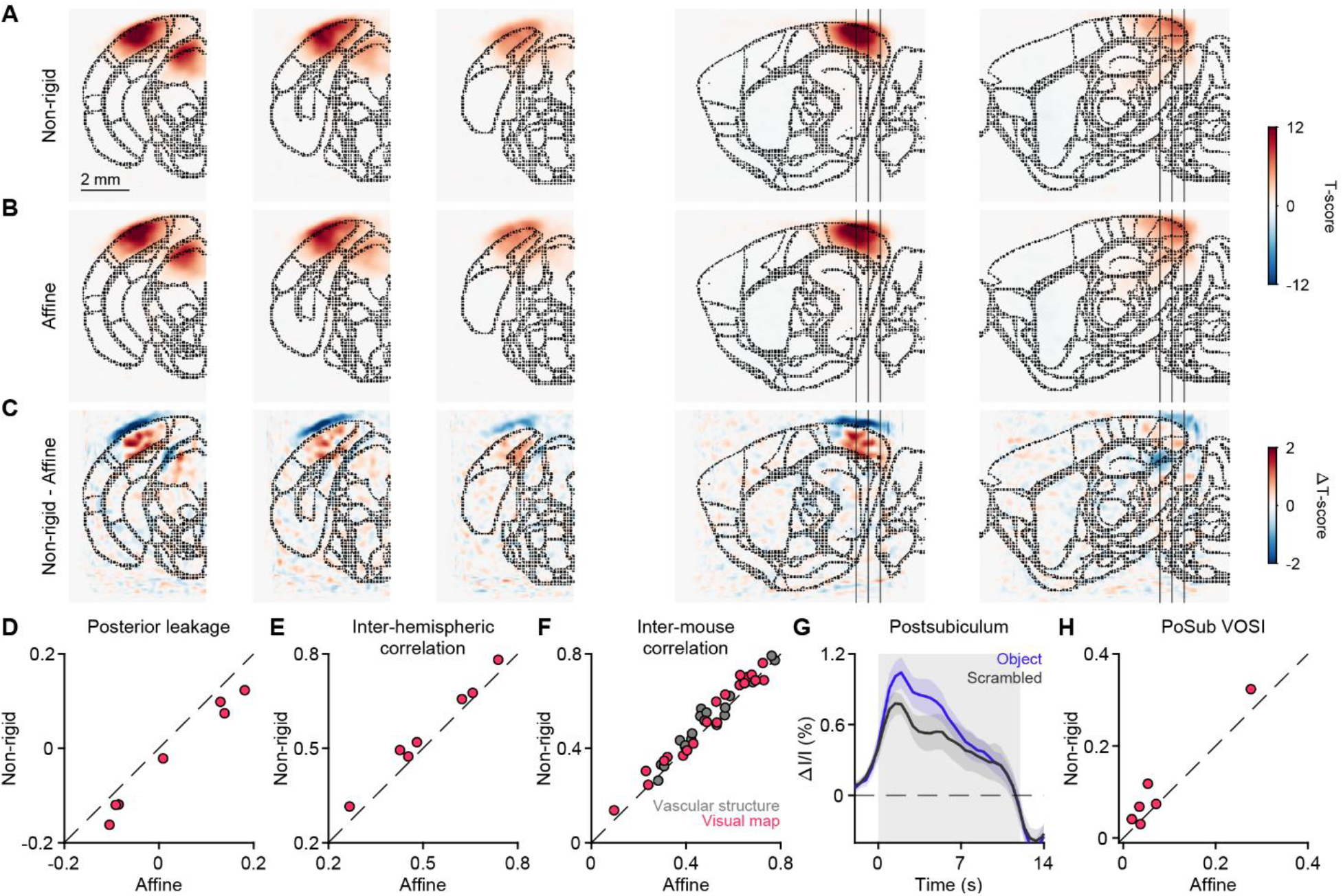
Quantitative evaluation of non-rigid *in vivo* fUS registration. **A**) Average t-score maps of visually evoked hemodynamic activity under non-rigid registration (mean across n=7 mice) across three coronal (columns 1–3) and two sagittal (columns 4–5) planes. Black dotted outlines indicate CCFv3 structural boundaries; external voxels are unmasked to assess boundary alignment. Vertical lines on sagittal slices mark the anteroposterior locations of the corresponding coronal sections. **B**) As in (A), for affine registration. **C**) Difference map between non-rigid and affine registrations (non-rigid minus affine). **D**) Posterior leakage index for positive t-score voxels, defined as (outside−inside)/(outside+inside) in the posterior quarter of the brain, comparing affine and non-rigid registration across mice (n=7 mice, p=0.016). **E)** Across-voxel correlation of t-score values between left and right hemispheres for responsive voxels within individual mice, comparing non-rigid and affine registration (n=7 mice, p=0.016). **F**) Pairwise inter-mouse correlation across registered volumes for anatomical vascular scans (gray) and visually evoked t-score maps (red), comparing non-rigid to affine registration. Both anatomical (n=21 pairs, p=0.004) and t-score volumes (p=0.004) show higher correlations for non-rigid compared to affine registration. **G**) Evoked relative signal intensity change in the postsubiculum in response to visual object (blue) and scrambled (black) stimuli over time under non-rigid registration. Shaded error bars denote SEM across mice (n=7); the gray shaded box indicates the visual stimulus window. **H**) Visual object sensitivity index (VOSI) in the postsubiculum is larger for non-rigid compared to affine registration (n=7 mice, p=0.047). Dashed lines in (D-F) and (H) denote identity; all statistical comparisons were performed through Wilcoxon signed-rank tests.

